# HIF1α-AS1 is a DNA:DNA:RNA triplex-forming lncRNA interacting with the HUSH complex

**DOI:** 10.1101/2021.10.11.463905

**Authors:** Matthias S. Leisegang, Jasleen Kaur Bains, Sandra Seredinski, James A. Oo, Nina M. Krause, Chao-Chung Kuo, Stefan Günther, Nevcin Sentürk Cetin, Timothy Warwick, Can Cao, Frederike Boos, Judit Izquierdo Ponce, Rebecca Bednarz, Chanil Valasarajan, Dominik Fuhrmann, Jens Preussner, Mario Looso, Soni S. Pullamsetti, Marcel H. Schulz, Flávia Rezende, Ralf Gilsbach, Beatrice Pflüger-Müller, Ilka Wittig, Ingrid Grummt, Teodora Ribarska, Ivan G. Costa, Harald Schwalbe, Ralf P. Brandes

## Abstract

DNA:DNA:RNA triplexes that are formed through Hoogsteen base-pairing have been observed *in vitro*, but the extent to which these interactions occur in cells and how they impact cellular functions remains elusive. Using a combination of bioinformatic techniques, RNA/DNA pulldown and biophysical studies, we set out to identify functionally important DNA:DNA:RNA triplex-forming long non-coding RNAs (lncRNA) in human endothelial cells. The lncRNA HIF1α-AS1 was retrieved as a top hit. Endogenous HIF1α-AS1 reduced the expression of numerous genes, including EPH Receptor A2 and Adrenomedullin through DNA:DNA:RNA triplex formation by acting as an adapter for the repressive human silencing hub complex (HUSH). Moreover, the oxygen-sensitive HIF1α-AS1 was down-regulated in pulmonary hypertension and loss-of-function approaches not only resulted in gene de-repression but also enhanced angiogenic capacity. As exemplified here with HIF1α-AS1, DNA:DNA:RNA triplex formation is a functionally important mechanism of trans-acting gene expression control.

## Introduction

Long non-coding RNAs (lncRNAs) represent the most diverse, plastic and poorly understood class of ncRNA^1^. Their gene regulatory mechanisms involve formation of RNA-protein, RNA-RNA or RNA-DNA complexes^1^. RNA-DNA interactions occur either in heteroduplex (DNA:RNA) or triplex strands (DNA:DNA:RNA). In triplexes, double-stranded DNA (dsDNA) accommodates the single-stranded RNA in its major groove^2^. The binding occurs via Hoogsteen or reverse Hoogsteen hydrogen bonds with a purine-rich sequence of DNA to which the RNA strand binds in a parallel or antiparallel manner. Hoogsteen bonds are weaker than Watson-Crick bonds, resulting in Hoogsteen pairing rules being more flexible^3^.

*Ex vivo* triplex formation relies on different biophysical methods including circular dichroism- (CD) and nuclear magnetic resonance-spectroscopy (NMR)^4–6^. Even with these techniques it can be challenging to discriminate DNA-RNA heteroduplexes from triplexes and analyses are usually restricted to oligonucleotides of a limited length. Nevertheless, a few lncRNAs have been suggested to form triplexes with dsDNA, however, triplex studies using living cells are still in early development^4, 6–13^. *In silico* analyses of RNA-DNA triplex formation predicted several genomic loci and lncRNAs to form triplexes^14^. In line with this, a global approach in HeLa S3 and U2OS cells to isolate triplex-forming RNAs on a genome-wide scale yielded several RNA:DNA triplex-forming lncRNAs^15^.

In addition to the sparse initial findings of triplex formation within cells, several other open questions remain: What is the physiological relevance of triplex-forming lncRNAs and are these cell- and tissue-type specific? What is the mechanism of action of triplex-forming lncRNAs? Do they disturb transcription in a similar way to R-loops^16^ or recruit certain protein complexes to DNA in a site-specific manner? Regarding the latter aspect, Polycomb Repressive Complex 2 (PRC2) has been identified as a target of the lncRNAs HOX Transcript Antisense RNA (HOTAIR), FOXF1 Adjacent Non-Coding Developmental Regulatory RNA (FENDRR) and Maternally Expressed 3 (MEG3)^4, 12, 13^, but, given the highly promiscuous nature of PRC2, this function remains controversial. Other examples of protein interactors involve e.g. E2F1 and p300, which are recruited by the triplex-forming antisense lncRNA KHPS1 to activate gene expression of the proto-oncogene sphingosine kinase 1 (SPHK1) in cis^7, 10^.

Much of today’s *in vivo* RNA research heavily relies on immortalized cell lines. Although such model systems are well suited for transfection or genomic manipulation, they are highly de-differentiated and exhibit reaction patterns such as unlimited growth and immortalization - characteristics not observed in primary cells^17^. Considering that lncRNAs are expressed in a species-, tissue- and differentiation-specific manner^1^, biological evidence for lncRNA functions in primary cells is limited.

Among such cells, endothelial cells stand out due to their well documented importance in regeneration, angiogenesis and tissue vascularization. Indeed, endothelial cell dysfunction is one of the main drivers of systemic diseases like diabetes and inflammation^18^.

Here, we combined molecular biology and biophysics, bioinformatics and physiology to systematically uncover the role of triplex-forming lncRNAs in endothelial cells. This approach identified HIF1α-AS1 as a *trans*-acting triplex-forming lncRNA that controls vascular gene expression in endothelial cells with implications for vascular disease.

## Results

### HIF1α-AS1 is a triplex-associated lncRNA

To identify triplex-associated lncRNAs, we used Triplex-Seq data from U2OS and HeLa S3 cells^15^. Triplex-Seq relies on the isolation of RNase H-resistant RNA-DNA complexes from cells followed by DNA- and RNA-Seq^15^. The data comprised all RNA entities and was filtered for lncRNAs, resulting in 989 (for HeLa S3, **Sup. Table 1**) and 1386 (for U2OS, **Sup. Table 2**) lncRNA regions associated with triplexes, with an overlap of 280 regions between the two cell lines (**Fig. 1a**). To further narrow down this set of enriched triplex-associated lncRNAs, parameters for specificity (fold enrichment >10, minus_log10(P) >20) were increased so that 11 lncRNA candidates with high confidence remained. Subsequently, these were correlated to Encode and FANTOM5 Cap Analysis of Gene Expression (CAGE)^19–21^ data. Of the 11 candidates, only 5 (RMRP, HIF1α-AS1, RP5-857K21.4, SCARNA2 and SNHG8) were expressed in endothelial cells. All 5 candidates were predicted as non-coding by the online tools Coding Potential Assessment Tool (CPAT) and coding potential calculator 2 (CPC2) and at least partially nuclear localized by Encode and FANTOM5 CAGE (**Fig. 1a**). To further analyze these candidates, the Triplex-Seq enriched regions were manually inspected in the IGV browser. This led to the exclusion of SNHG8 as the triplex-associated regions within this lncRNA were exclusively within the overlapping small nucleolar RNA 24 (SNORA24) gene. In the case of the other candidates, triplex- association was within the individual lncRNA gene body. The cumulative fold enrichment of the remaining lncRNAs in the Triplex-Seq dataset illustrated strong triplex-association (**Extended data Fig. 1a**). To verify the candidates experimentally, RNA immunoprecipitation (RIP) with antibodies against dsDNA and with or without RNase H treatment in human endothelial cells was performed. RNase H, which cleaves the RNA in DNA-RNA heteroduplexes (R-loops)^22^, revealed that HIF1α-AS1 was the strongest triplex-associated lncRNA (**Fig. 1b**).

**Fig. 1:**
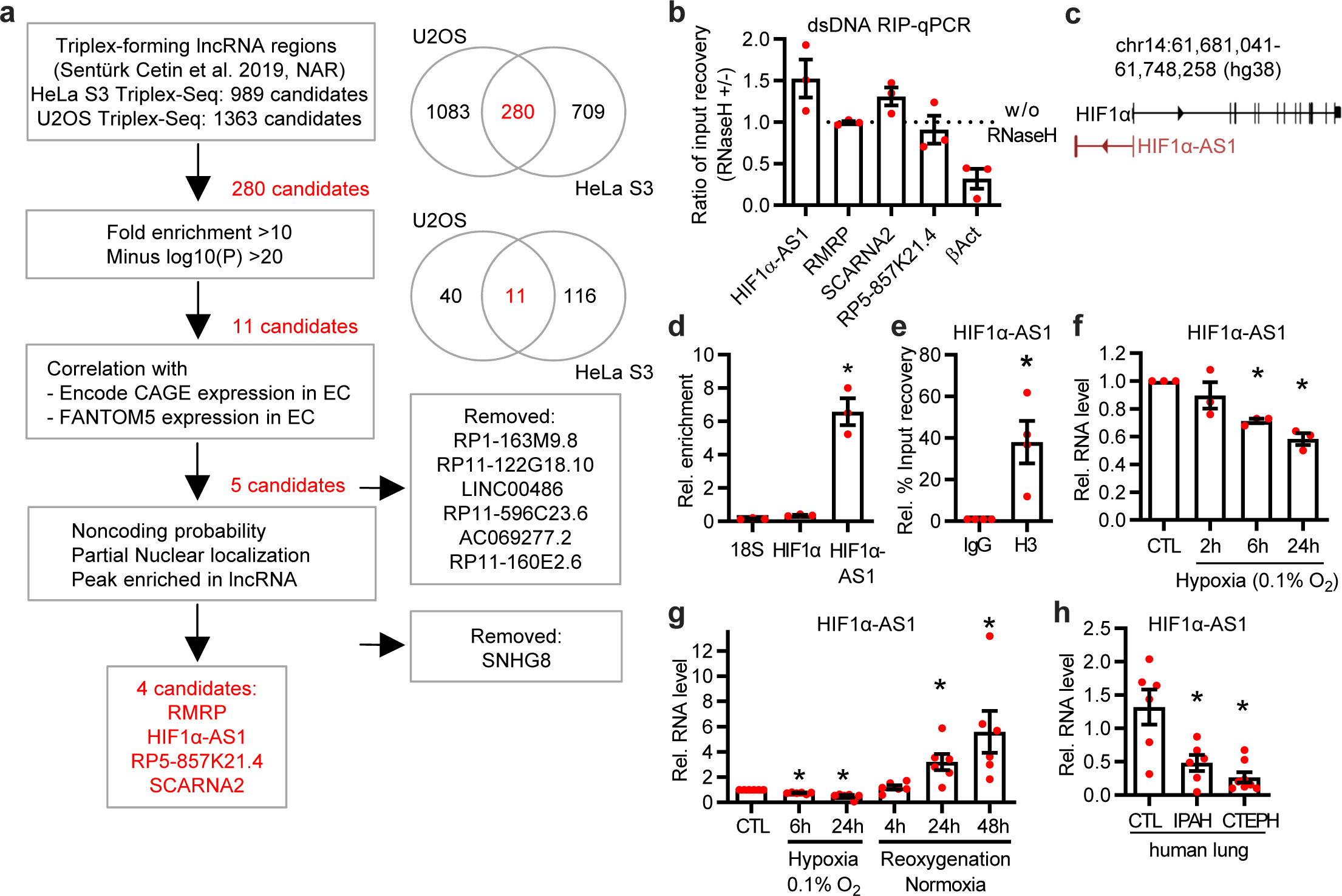
HIF1α-AS1 is a triplex- and DNA-associated RNase H-insensitive lncRNA in endothelial cells. **a**, Overview of the identification of endothelial-expressed triplex-forming lncRNAs. LncRNAs from a previous Triplex-Seq study in HeLa S3 and U2OS were overlapped, filtered with high stringency and analyzed for nuclear expression in endothelial cells with Encode and FANTOM5 CAGE data followed by analyses for noncoding probability and enriched peaks in the Triplex-Seq data. **b**, RNA- immunoprecipitation with anti-dsDNA followed by qPCR (RIP-qPCR) targeting the lncRNA candidates in HUVEC. Samples were treated with or without RNase H. βAct served as control for RNase H- mediated degradation. n=3. **c**, Scheme of the human genomic locus of HIF1α-AS1. **d**, RT-qPCR after anti-dsDNA-RIP in HUVEC. HIF1α and 18S rRNA served as negative control. One-way ANOVA with Tukey’s post hoc test, n=3. **e**, RIP-qPCR with anti-histone3 (H3) in HUVEC. Data was normalized against GAPDH. Paired t-test, n=4. **f**, RT-qPCR of HIF1α-AS1 in HUVEC treated with hypoxia (0.1% O2) for the indicated time points. Normoxia served as negative control (CTL). n=3, One-Way ANOVA with Bonferroni post hoc test. **g**, RT-qPCR of HIF1α-AS1 in HUVECs treated with hypoxia (0.1% O2) followed by reoxygenation with normoxia (after 24 h hypoxia) for the indicated time points. n=6, One-Way ANOVA with Dunnett’s post hoc test. **h**, RT-qPCR of HIF1α-AS1 in lungs from control donors (CTL, n=6) or patients with idiopathic pulmonary arterial hypertension (IPAH, n=6) or chronic thromboembolic pulmonary hypertension (CTEPH, n=8). One-Way ANOVA with Tukey’s post hoc test. Error bars are defined as mean +/- SEM. *p<0.05.

**Table 1.**
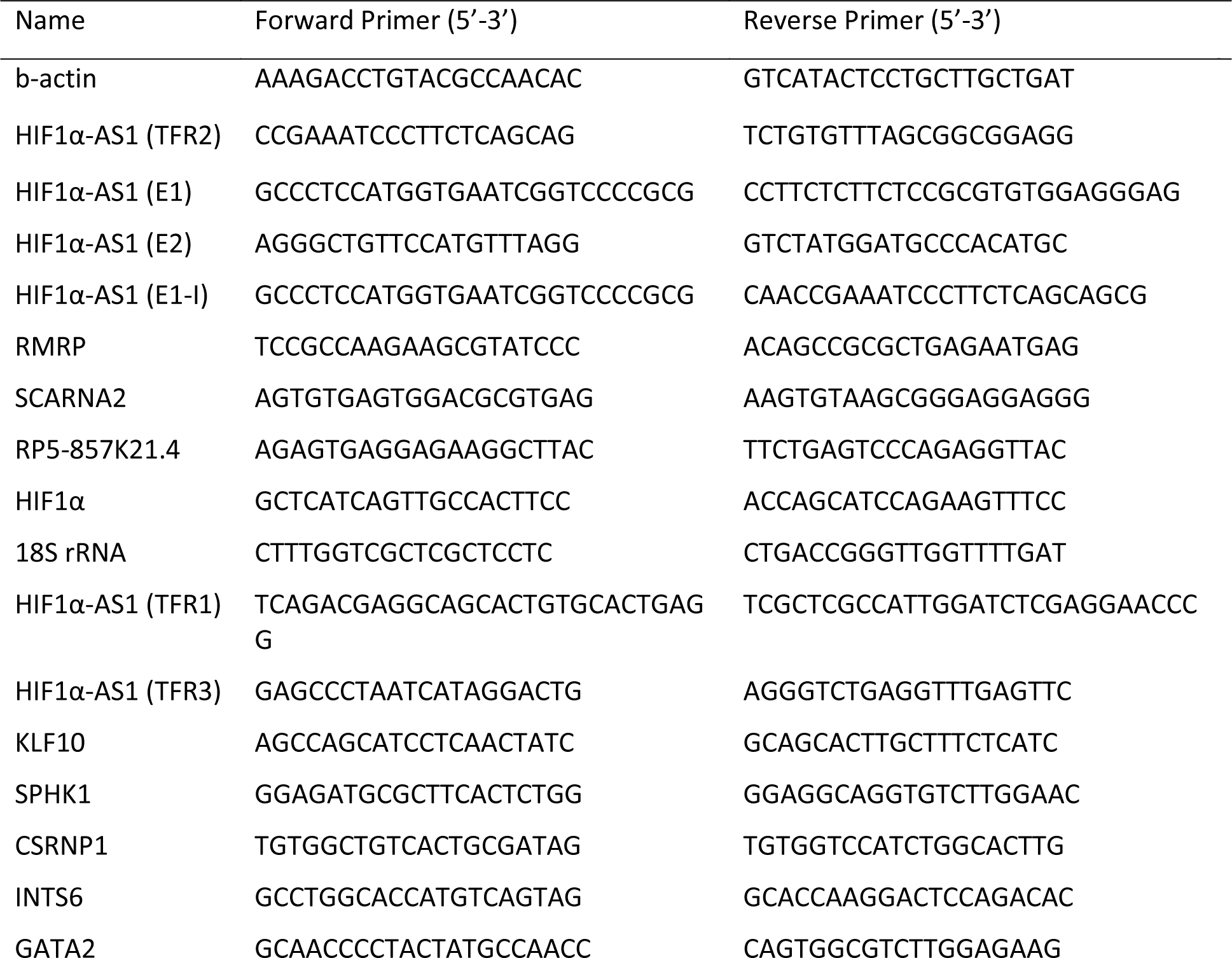
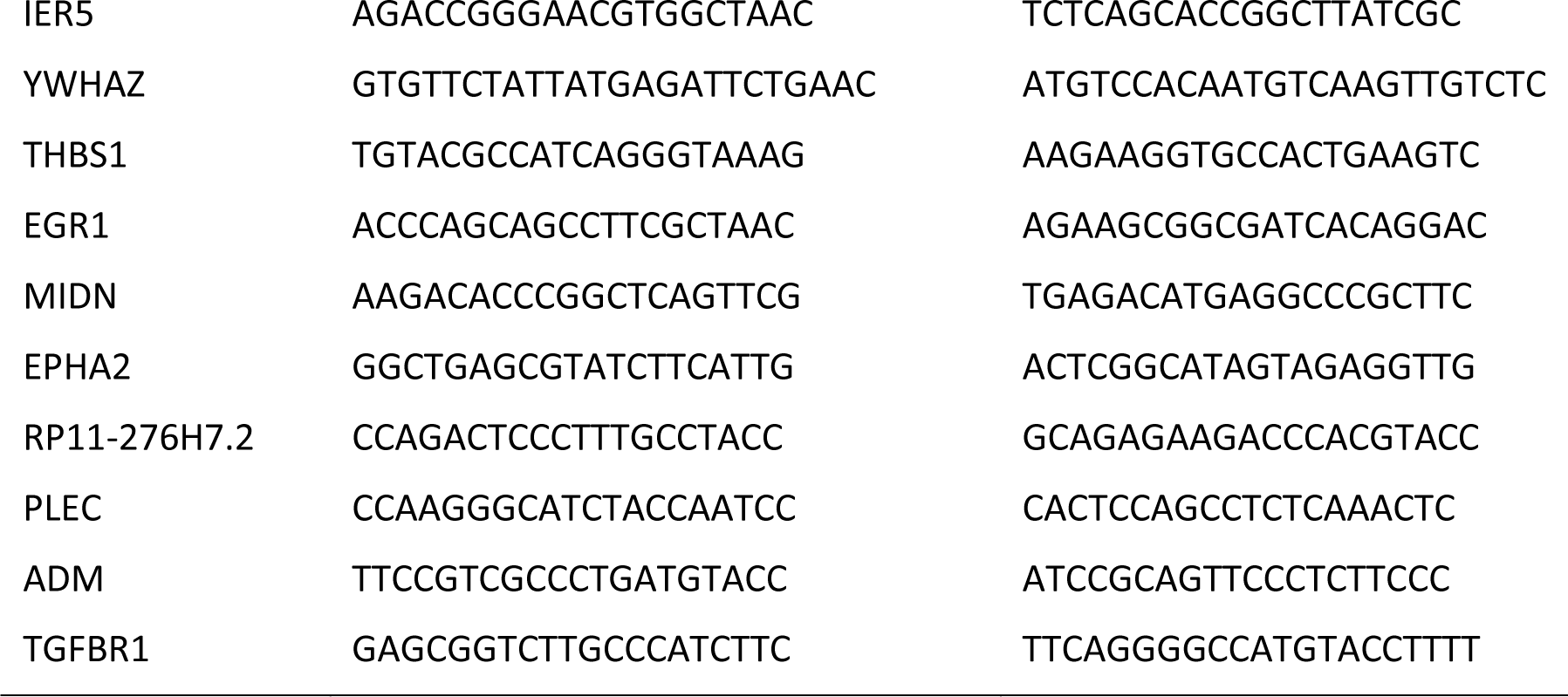
List of primers for qRT-PCR.

**Table 2.**
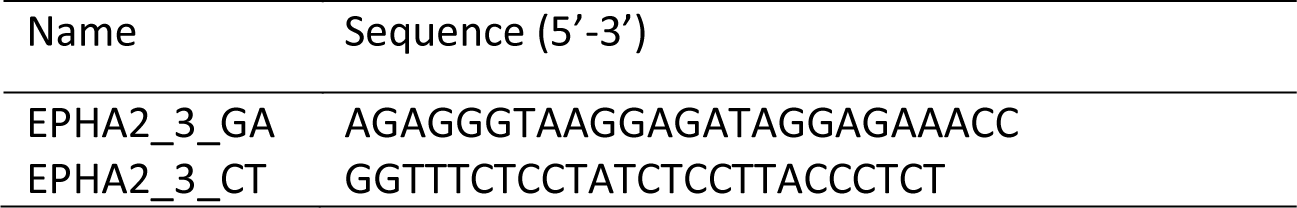
DNA oligos used for triplex target sites.

Genomically, HIF1α-AS1 is located on the antisense strand of the Hypoxia-inducible factor 1α gene (**Fig. 1c**). The lncRNA was specifically enriched in nuclear DNA, whereas HIF1α mRNA and 18S rRNA were not (**Fig. 1d**). Moreover, RIP with anti-histone 3 (**Fig. 1e**) indicated that HIF1α-AS1 is bound to dsDNA in the chromatin environment.

### HIF1α-AS1 is disease-relevant

Only a few studies have so far reported the biological relevance of HIF1α-AS1. Increased HIF1α-AS1 expression has been reported in thoracoabdominal aortic aneurysms^23^. HIF1α-AS1 was also suggested as a biomarker in colorectal carcinoma^24^. Functionally, HIF1α-AS1 is pro-apoptotic and anti-proliferative in vascular smooth muscle, Kupffer and umbilical vein endothelial cells^25–27^.

As HIF1α is a central regulator of oxygen-dependent gene expression^18^, we decided to measure the expression of HIF1α-AS1 in endothelial cells in altered oxygen and disease conditions. Hypoxia led to a decrease in HIF1α-AS1 expression in endothelial and pulmonary artery smooth muscle cells (paSMC) (**Fig. 1f**, **Extended data Fig. 1b**), which was restored in endothelial cells after 4 h and even surpassed basal levels after 24 h of normoxic conditions (**Fig. 1g**). Importantly, HIF1α-AS1 was downregulated in endothelial cells isolated from human glioblastoma (**Extended data Fig. 1c**) and in lungs from patients with end stage idiopathic pulmonary arterial hypertension (IPAH) or chronic thromboembolic pulmonary hypertension (CTEPH) (**Fig. 1h**). In paSMCs isolated from pulmonary arteries of patients with IPAH, HIF1α-AS1 was strongly decreased (**Extended data Fig. 1d**). Together, these data demonstrate that HIF1α-AS1 is an oxygen-dependent and disease-relevant lncRNA.

### HIF1α-AS1-triplex binding suppresses target gene expression

Triplex-Seq provides evidence for existing triplex forming regions of the RNA (TFR) and triplex target sites (TTS) within the DNA but the details of exactly which TFR and TTS interact cannot be derived from Triplex-Seq. To identify the TFRs within HIF1α-AS1 as well as HIF1α-AS1-dependent TTS, a combination of bioinformatics and wet lab approaches were used: An Assay for Transposase-Accessible Chromatin with high-throughput sequencing (ATAC-Seq) was performed after HIF1α-AS1 knockdown to identify DNA target sites in human endothelial cells. LNA-GapmeRs targeting HIF1α- AS1 led to a strong knockdown of the lncRNA (**Extended data Fig. 1e**). Triplex Domain Finder (TDF) predicted the TFRs within HIF1α-AS1 to target DNA regions around genes that displayed altered ATAC-Seq peaks after HIF1α-AS1 silencing (**Fig. 2a**). The software identified three statistically significant TFRs (TFR1-3) within the pre-processed HIF1α-AS1 RNA (**Fig. 2b**). There was also a high incidence of triplex-prone motifs predicted in regions whose chromatin state was altered in the ATAC-Seq data after HIF1α-AS1 knockdown (**Fig. 2c**, **Sup. Tables 3-5**). Of these TTS, 38 overlapped within all three TFRs (**Fig. 2d**). To identify which TFR is most strongly associated with triplexes, RIP with S9.6 antibodies recognizing RNA-DNA association was performed. RNA-DNA associations remaining after RNase H treatment excluded the possibility that these were RNA-DNA heteroduplexes. Of the three HIF1α-AS1 TFRs, TFR2 was identified as the TFR most resistant to RNase H (**Fig. 2e**). TFR2 is located intronically 478 nucleotides (nt) downstream of Exon1 and was detected by RT-PCR within nuclear isolated RNA with primers covering the first 714 nt (E1-I) of the pre- processed HIF1α-AS1 (**Extended data Fig. 1f**). Triplex-prone motifs in their target regions yielded more than 20 different associated genes, some of which displayed a high number of DNA binding sites (**Fig. 2f**). If this binding of the lncRNA is truly relevant for the individual target gene, then a change in target gene expression would be expected. Importantly, in response to the downregulation of HIF1α-AS1 with LNA-GapmeRs the expression of the following triplex target genes increased: ADM, PLEC, RP11-276H7.2, EPHA2, MIDN and EGR1 (**Fig. 2g**). Interestingly, as exemplified by the target genes HIF1α, EPHA2 and ADM, the triplex target sites are often located close to the 5’ end of the gene. In this region histone modifications, transcription factor binding and chromatin conformation often have the greatest effect on promoter function and gene expression (**Fig. 2h**).

**Fig. 2:**
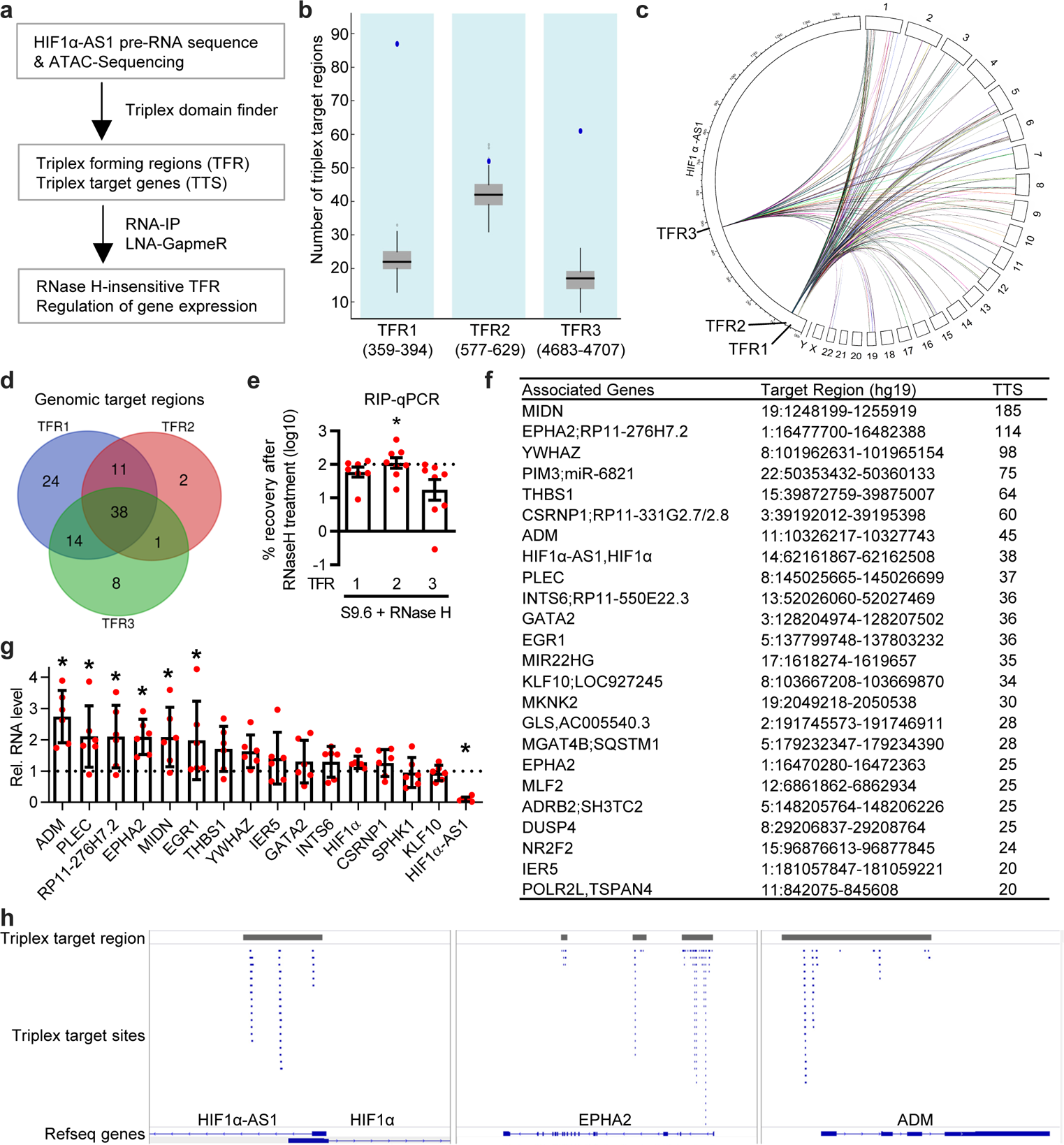
HIF1α-AS1 potentially forms DNA:DNA:RNA triplexes. **a**, Overview of the identification of HIF1α-AS1 triplex forming regions (TFR) and their DNA triplex target sites (TTS) with triplex domain finder (TDF). HIF1α-AS1 pre-RNA and ATAC-Seq of HUVECs treated with or without LNA GapmeRs against HIF1α-AS1 were used as input. RIP and LNA GapmeRs were used to validate the findings obtained by TDF. **b**, Number of triplex target regions of three statistically significant TFRs of HIF1α- AS1 identified with TDF. Numbers in brackets represent the position of the individual TFR within HIF1α-AS1 pre-RNA. All TFRs have a significantly higher number of triplex target regions in targets (blue) than non-target regions (grey). **c**, Circos plot showing the localization of the individual TFR within HIF1α-AS1 pre-RNA and its interaction with the chromosomal TTS. **d**, Overlap of TTS of the individual TFRs of HIF1α-AS1. **e**, Identification of RNase H-resistent TFRs. RIP with S9.6 RNA/DNA hybrid antibody with or without RNase H treatment in HUVEC followed by qPCR for the TFRs. Ratio of %-input recovery with/without RNase H treatment is shown. n=8, paired t-test. **f**, HIF1α-AS1 TFR2 top target genes, their genomic location and number of TTS identified by TDF. **g**, RT-qPCR of triplex target genes of TFR2 after knockdown of HIF1α-AS1 in HUVEC. n=6, One-Way ANOVA with Holm‘s Sidak post hoc test. **h**, Three different triplex target regions of HIF1α-AS1 are shown. Triplex target regions are highlighted in grey, triplex target sites are shown in blue. Error bars are defined as mean +/- SEM. *p<0.05.

These data indicate that HIF1α-AS1 contains triplex forming regions and target sites important for the regulation of gene expression.

### HIF1α-AS1 TFR2 RNA forms triplexes with EPHA2 and ADM

Our analysis identified HIF1α-AS1 TFR2 as the best suited candidate for verification of triplex formation of the lncRNA using biophysical and biochemical techniques. To monitor triplex formation of HIF1α-AS1, EPHA2 was chosen as the target gene due to its abundance of triplex target sites (**Fig. 2f, Fig. 2h**), its regulatory potential (**Fig. 2g**) and its importance for vascularization^28^. The formation of DNA:DNA:RNA triplexes between lncRNA HIF1α-AS1 TFR2 and its proposed DNA target site within intron 1 of EPHA2 was characterized by solution NMR spectroscopy, electrophoretic mobility shift assay (EMSA) and CD-spectroscopy. ^1^H-1D NMR spectra were recorded for EPHA2 DNA duplex, HIF1α-AS1 TFR2 RNA (TFO2-23), EPHA2:HIF1α-AS1_TFR2 heteroduplex and EPHA2:HIF1α-AS1_TFR2 triplex at different temperatures. Using 10 eq HIF1α-AS1 TFR2 RNA, triplex ^1^H NMR imino signals were observed in a spectral region between 9 and 12 ppm providing further evidence that HIF1α-AS1 was associated with EPHA2 through Hoogsteen base pairing (**Fig. 3a**). Moreover, HIF1α-AS1 TFR2 RNA formed a low mobility DNA–RNA complex with the radiolabeled EPHA2 DNA target sequence in electrophoretic mobility shift assays (EMSA). The shift in mobility retardation was dependent on the TFR2 transcript length (**Fig. 3b**). We also used CD-spectroscopy to confirm triplex formation of HIF1α- AS1 TFR2 on EPHA2. The CD spectrum indicated typical features for triplex formation, such as a positive small peak at ∼220 nm, two negative peaks at ∼210 nm and ∼240 nm and a blue-shift of the peak at ∼270 nm, which was distinct from the EPHA2 DNA duplex or the heteroduplex spectra (**Fig. 3c**). This confirmed the existence of EPHA2:HIF1α-AS1 TFR2 triplexes. Additionally, we performed UV melting assays and obtained melting temperatures Tm (RNA-DNA heteroduplex) = 53.48 ± 0.32 °C, Tm (DNA-DNA duplex) = 70.73 ± 0.22 °C and Tm (DNA-DNA-RNA triplex) = 54.17 ± 0.23 °C with a very broad second melting point around 70 °C. The biphasic melting transition is a distinct feature of triplex formation, where the first melting temperature corresponds to melting of Hoogsteen hydrogen bonds that stabilize the triplex and the second for the melting of the Watson-Crick base pairing at higher temperatures (**Fig. 3d**).

**Fig. 3:**
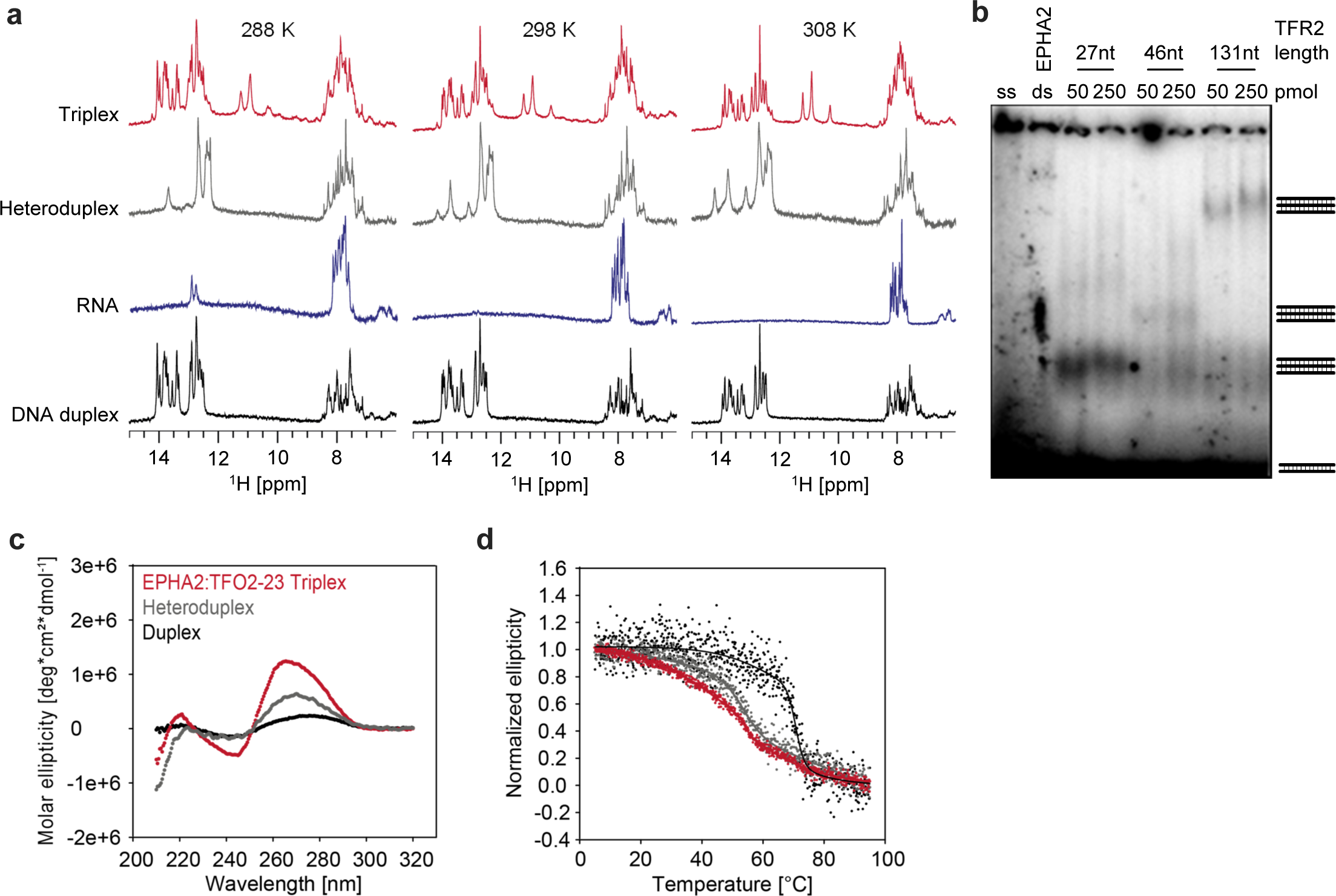
HIF1α-AS1 TFR2 RNA forms *in vitro* DNA:DNA:RNA triplexes with the predicted DNA target region in EPHA2. **a**, ^1^H-1D NMR spectra of the EPHA2 DNA duplex (black), HIF1α-AS1 TFR2 RNA (blue), heteroduplex (dark grey) and EPHA2:HIF1α-AS1-TFR2 triplex (red) in a temperature range between 288-308 K. **b**, Electromobility shift assay of EPHA2 ssDNA or dsDNA (ss or ds) alone or the dsDNA in combination with HIF1α-AS1-TFR2. Two different RNA dosages (50 or 250 pmol) and three different HIF1α-AS1-TFR2 RNA lengths (27 nt, 46 nt, 131 nt) were used **c**, Circular dichroism spectra of the EPHA2 DNA duplex (black), the heteroduplex (dark grey) and EPHA2:HIF1α-AS1-TFR2 triplex (red) measured at 298 K. **d**, UV melting assay of the EPHA2 DNA duplex (black), the heteroduplex (dark grey) and EPHA2:HIF1α-AS1-TFR2 triplex (red).

To confirm the formation of triplexes with lower equivalents, stabilized triplex formation was investigated: the intermolecular dsDNA form from two complementary antiparallel DNA strands was changed into a hairpin construct, where both DNA strands were linked with a 5 nt thymidine-linker and duplex formation thus became intramolecular. With this approach, triplex formation was obtained with 3 eq RNA, indicating that triplex formation is favored under those conditions as expected. ^1^H-1D NMR spectra of hairpin EPHA2_CTGA and 15N HIF1α-AS1 TFR2:EPHA_CTGA triplex indicated changes in the Hoogsteen region (9-12 ppm) and the spectral region of imino (12-14 ppm) and amino signals (7-8.5 ppm) (**Extended data Fig. 2a**). In addition to EPHA2, we also tested ADM, a preprohormone involved in endothelial cell function^29^. For ADM_CTGA:HIF1α-AS1 TFR2 triplex, the new imino protons in the Hoogsteen region arose at lower temperatures (**Extended data Fig. 2b**). For both ADM_CTGA and EPHA2_CTGA triplex constructs the CD spectra showed an increased negative ellipticity at ∼240 nm and positive ellipticity at ∼270 nm (**Extended data Fig. 2c,e**). Further, the UV melting data verified the triplex stabilization with higher melting temperatures and defined melting transitions upon DNA hairpin formation. For the EPHA2_CTGA:HIF1α-AS1 TFR2 (TFO2-23) triplex we obtained a first melting point at Tm (1^st^ triplex) = 50.08 ± 0.51 °C, a second melting point Tm (2^nd^ triplex) = 79.90 ± 0.10 °C and Tm (DNA hairpin) = 80.41 ± 0.10 °C (**Extended data Fig. 2d**). The melting temperature of ADM DNA duplex Tm (DNA-DNA duplex) = 63.80 ± 0.20 °C increased for the ADM_CTGA hairpin Tm (DNA hairpin) = 95.76 ± 16.69 °C. For the ADM_CTGA:HIF1α-AS1 TFR2 (TFO2-23), we obtained a first melting point Tm (1st triplex) = 51.19 ± 0.68 °C and a second Tm (2nd triplex) = 82.86 ± 0.21 °C (**Extended data Fig. 2f**). The data demonstrate that HIF1α-AS1 TFR2 forms triplexes with EPHA2 and ADM dsDNA under regular and triplex-stabilized conditions upon DNA hairpin formation.

### TFR2 represses EPHA2 and ADM gene expression

The current data indicates that HIF1α-AS1 forms triplexes with EPHA2 and ADM, however, the mechanistic and functional consequences of this phenomenon are unclear. To investigate these aspects, gain and loss of function approaches were performed. Increasing the expression of HIF1α- AS1 using a dCas9-VP64 CRISPR activation system (CRISPRa) reduced the expression of EPHA2 and ADM (**Fig. 4a**). Conversely, downregulation of HIF1α-AS1 with a dCas9-KRAB repression system (CRISPRi) increased the expression of EPHA2 and ADM (**Fig. 4b**). Consistent with HIF1α-AS1 repressing EPHA2 and ADM gene expression, EPHA2 levels increased after knockdown of HIF1α-AS1 (**Fig. 2g**, **Fig. 4c**). EPHA2 has a multi-faceted role in angiogenesis^28, 30, 31^. In HUVEC, knockdown of EPHA2 with siRNAs strongly reduced its RNA and protein expression and inhibited angiogenic sprouting (**Fig. 4d&e**, **Extended data Fig. 3a-c**). Conversely, a knockdown of HIF1α-AS1 with LNA-GapmeRs increased basal, VEGF-A- and bFGF-mediated angiogenic sprouting (**Fig. 4f-g**, **Extended data Fig. 3d**), confirming the repressive effect of HIF1α-AS1 on EPHA2. To demonstrate directly that TFR2 is responsible for the regulation of EPHA2, we replaced TFR2 by genome editing using a recombinant Cas9-eGFP, a gRNA targeting TFR2 and different single-stranded oligodeoxynucleotides (ssODN) harboring either the published MEG3 TFR^4^ or a luciferase control sequence (**Fig. 4h**). Replacement of the TFR2 with the MEG3 TFR, which served as a positive control for a functional TFR repressing TGFBR1 expression^4^, yielded a reduction in TGFBR1 levels compared to the luciferase control (**Fig. 4i**). More importantly, the loss of TFR2 consequently led to a loss of HIF1α-AS1 TFR2, an upregulation of EPHA2 and partially of ADM (**Fig. 4j&k**, **Extended data Fig.3e**). These data demonstrate that TFR2 represses EPHA2 and ADM gene expression.

**Fig. 4:**
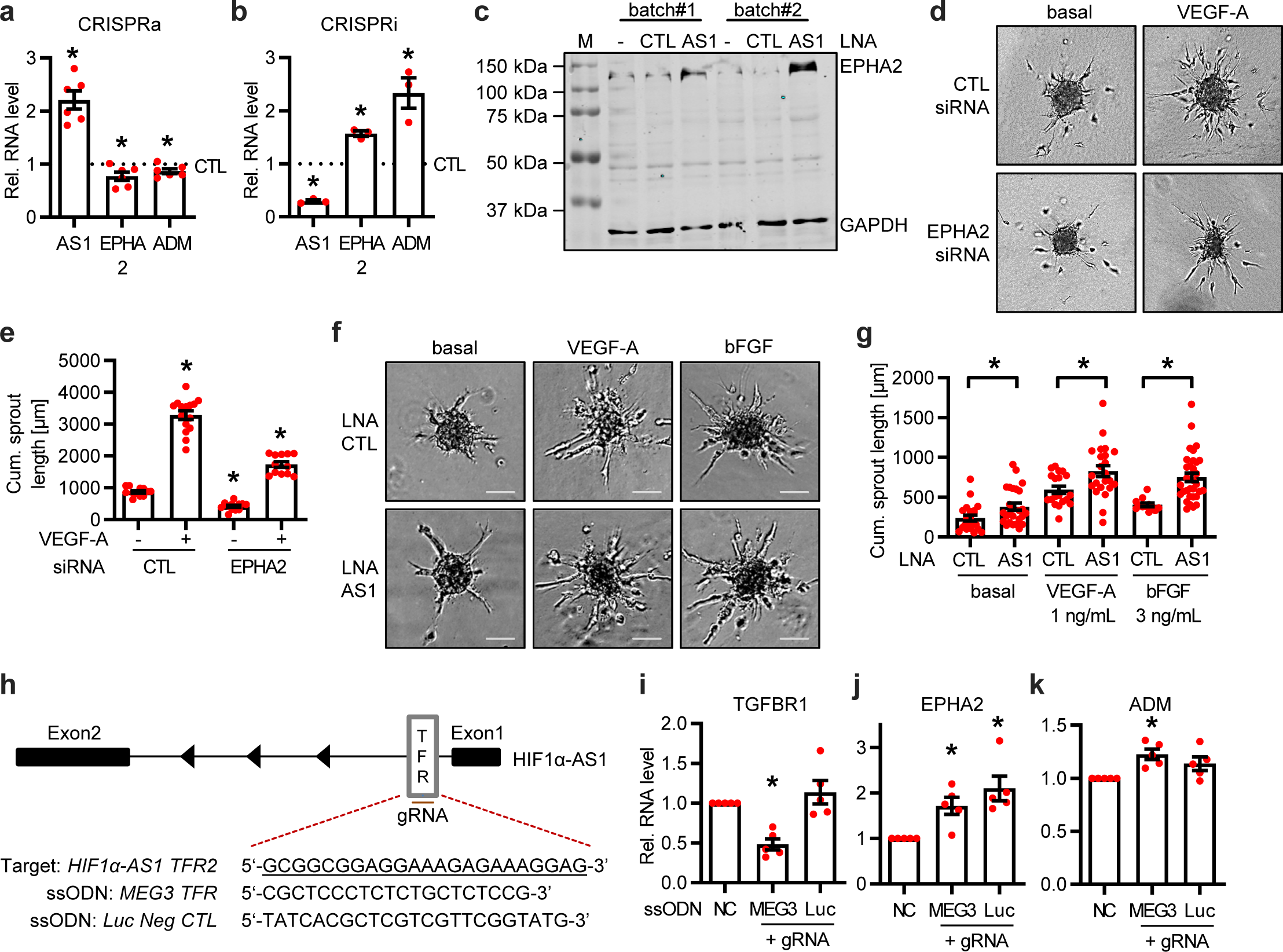
HIF1α-AS1 limits EPHA2 and ADM expression through TFR2. **a&b**, CRISPRa (a, n=6) or CRISPRi (b, n=3) targeting HIF1α-AS1 in HUVECs followed by RT-qPCR for HIF1α-AS1, EPHA2 and ADM. n=6, Paired t-test. **c**, Western blot with (AS1) or without (- and CTL) LNA GapmeR-mediated knockdown of HIF1α-AS1 in two different batches of HUVEC. GAPDH was used as loading control. M, marker. **d**, Spheroid outgrowth assay of HUVECs treated with or without siRNAs against EPHA2. Cells treated under basal or VEGF-A (1 ng/mL) conditions for 16 h are shown. **e**, Quantification of the cumulative sprout length from the spheroid assay seen in Fig. 4d. One-Way ANOVA with Bonferroni post hoc test. n=12-15. **f**, Spheroid outgrowth assay of HUVECs treated with LNA GapmeRs targeting HIF1α- AS1. Cells treated under basal, VEGF-A (1 ng/mL) or bFGF (3 ng/mL) conditions for 16 h are shown. LNA CTL served as negative control. Scale bar indicates 200 µm. **g**, Quantification of the cumulative sprout length from the spheroid outgrowth assay seen in Fig. 4f. One-Way ANOVA with Bonferroni post hoc test. n=12-32. **h**, Scheme of the CRISPR Arcitect approach. TFR2 of HIF1α-AS1 (underlined) was targeted with Cas9/gRNA and replaced with ssODNs including MEG3 TFR or a DNA fragment of luciferase negative control. **i-k**, RT-qPCR of TGFBR1 (i), EPHA2 (j) or ADM (k) after replacement of HIF1α-AS1-TFR2 with MEG3-TFR or a DNA fragment of a luciferase negative control. NC, nontemplate control. n=5, Paired t-test. Error bars are defined as mean +/- SEM. *p<0.05. AS1, HIF1α-AS1.

### HIF1α-AS1 binds to and recruits HUSH to triplex targets

To elucidate the mechanism by which HIF1α-AS1 represses gene expression, HIF1α-AS1-associated proteins were studied using RNA pulldown experiments. 3’biotinylated spliced HIF1α-AS1 lncRNA or 3’biotinylated pcDNA3.1+ negative control were incubated in nuclear extracts from HUVECs and RNA-associated proteins were identified by electrospray ionization mass spectrometry, which retrieved M-phase phosphoprotein 8 (MPP8)-a component of the human silencing hub (HUSH) complex- as top hit (**Fig. 5a-b**, **Sup. Table 6**). The HUSH-complex is a nuclear machinery originally thought to mediate gene silencing during viral infection by recruiting the SET Domain Bifurcated Histone Lysine Methyltransferase 1 (SETDB1) which methylates H3K9^32^. The HUSH complex has not yet been studied in vascular cells and an interaction of its core protein MPP8 with lncRNAs has not been reported. To support our finding, RIP revealed that HIF1α-AS1 and its TFR2, but not HIF1α mRNA, interact with MPP8 (**Fig. 5c**, **Extended data Fig. 4a-b**). Furthermore, HIF1α-AS1 was highly enriched with H3K9me3 (**Fig. 5d**).

**Fig. 5:**
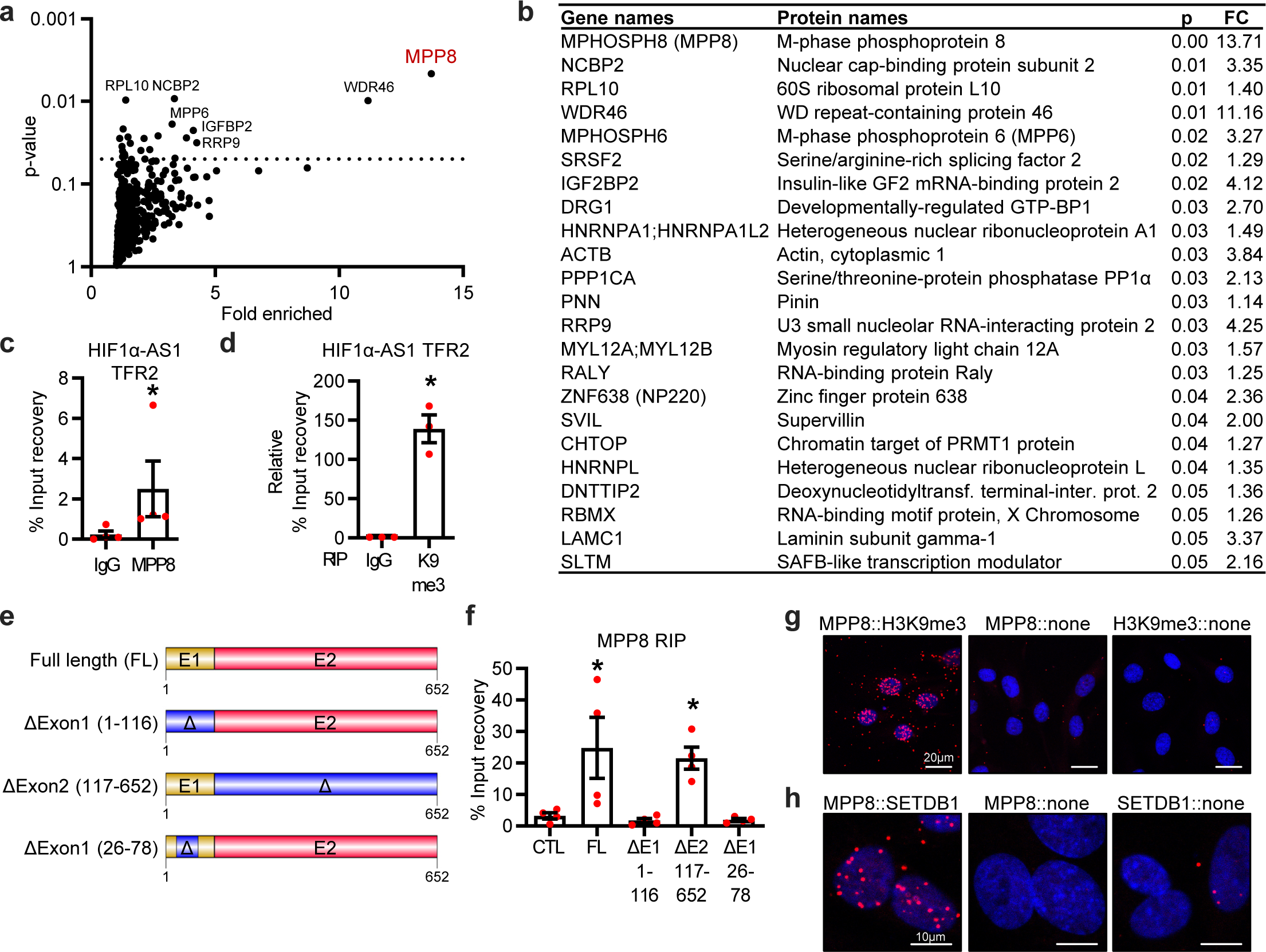
HIF1α-AS1 interacts directly with the HUSH complex member MPP8. **a**, Volcano plot of HIF1α-AS1 protein interaction partners after RNA pulldown assay and ESI-MS/MS measurements with fold enrichment and p-value. n=5. Proteins above the line (p<0.05) indicate significantly associated proteins. **b**, List of proteins enriched after RNA pulldown assay, their p-value and fold change. **c**, RIP with MPP8 antibodies and qPCR for HIF1α-AS1 TFR2. IgG served as negative control. n=4, Mann Whitney t-test. **d**, RIP with histone3-lysine9-trimethylation antibodies and qPCR for HIF1α-AS1 TFR2. IgG served as negative control. n=3, One-Way ANOVA with Dunnett‘s post hoc test. **e**, Scheme of the different HIF1α-AS1 RNAs used for *in vitro* RNA immunoprecipitation. **f**, RT-qPCR after *in vitro* binding assay of purified MPP8 with *in vitro* transcribed HIF1α-AS1 RNAs. MPP8 antibodies were used for RNA immunoprecipitation. An T7-MCS *in vitro* transcribed RNA served as negative control (CTL). FL, full length; E1, Exon1; E2, Exon2. Δ indicates the deleted nt from HIF1α- AS1 full length. **g-h**, Proximity ligation assay of HUVECs with antibodies against MPP8 and H3K9me3 (g) or MPP8 and SETDB1 (h). The individual antibody alone served as negative control. Red dots indicate polymerase amplified interaction signals. Scale bar indicates 20 µm (g) or 10 µm (h). Error bars are defined as mean +/- SEM. *p<0.05.

To map the RNA binding region of MPP8 on HIF1α-AS1, we used *cat*RAPID fragments^33^, an algorithm involving division of polypeptide and nucleotide sequences into fragments to estimate the interaction propensity of protein-RNA pairs. This highlighted potential binding regions within Exon1 (**Extended data 4c**). To substantiate these data experimentally, *ex vivo* bindings assays were performed between fragments of HIF1α-AS1 and recombinant MPP8 (**Fig. 5e**). MPP8 interacted directly with HIF1α-AS1 full length and a HIF1α-AS1 mutant lacking Exon2 (**Fig. 5f**). In contrast and in accordance with the *cat*RAPID prediction, deletion of Exon1 (nucleotides 26-78nt in particular) prevented the interaction (**Fig. 5f**), indicating that this region of HIF1α-AS1 is critical for the interaction of HIF1α-AS1 with MPP8.

To demonstrate that HIF1α-AS1 acts through HUSH complex recruitment, we first tested whether this complex exists in endothelial cells. Proximity ligation assays with antibodies against MPP8, dsDNA, H3K9me3 and SETDB1 confirmed the association of MPP8 with dsDNA (**Extended data Fig. 4d**), H3K9me3 (**Fig. 5g**) and SETDB1 (**Fig. 5h**) in the nuclei of endothelial cells, indicating that the complex is present in endothelial chromatin.

Chromatin immunoprecipitation (ChIP) with and without RNase A revealed that targeting of MPP8 to the HIF1α-AS1 TTS of EPHA2 and ADM was attenuated after RNA depletion (**Fig. 6a**). To demonstrate the dependence of the interactions with the TTS on HIF1α-AS1, ChIP experiments with antibodies targeting SETDB1, MPP8 and NP220 with or without knockdown of HIF1α-AS1 were performed. NP220 (ZNF638), which is another member of the HUSH complex, interacted with HIF1α-AS1, albeit to a lower degree than MPP8 (**Fig. 5b**). The binding of SETDB1 and MPP8, but not of NP220, to the triplex target sites of HIF1α-AS1 required the presence of the lncRNA (**Fig. 6b-c**) suggesting that these interactions facilitate epigenetic processes and ultimately regulate gene expression. ATAC-Seq confirmed that these factors act in the region of the TTS: After knockdown of HIF1α-AS1, SETDB1 or MPP8, the chromatin accessibility of both the EPHA2 and ADM transcriptional start sites were reduced. An increase in accessibility to the region downstream of the EPHA2 TTS was detected (**Fig. 6d**). These data indicate that the triplex formation by HIF1α-AS1 is important for fine-tuning chromatin accessibility locally and thereby gene expression of EPHA2 and ADM through SETDB1 and MPP8.

**Fig. 6:**
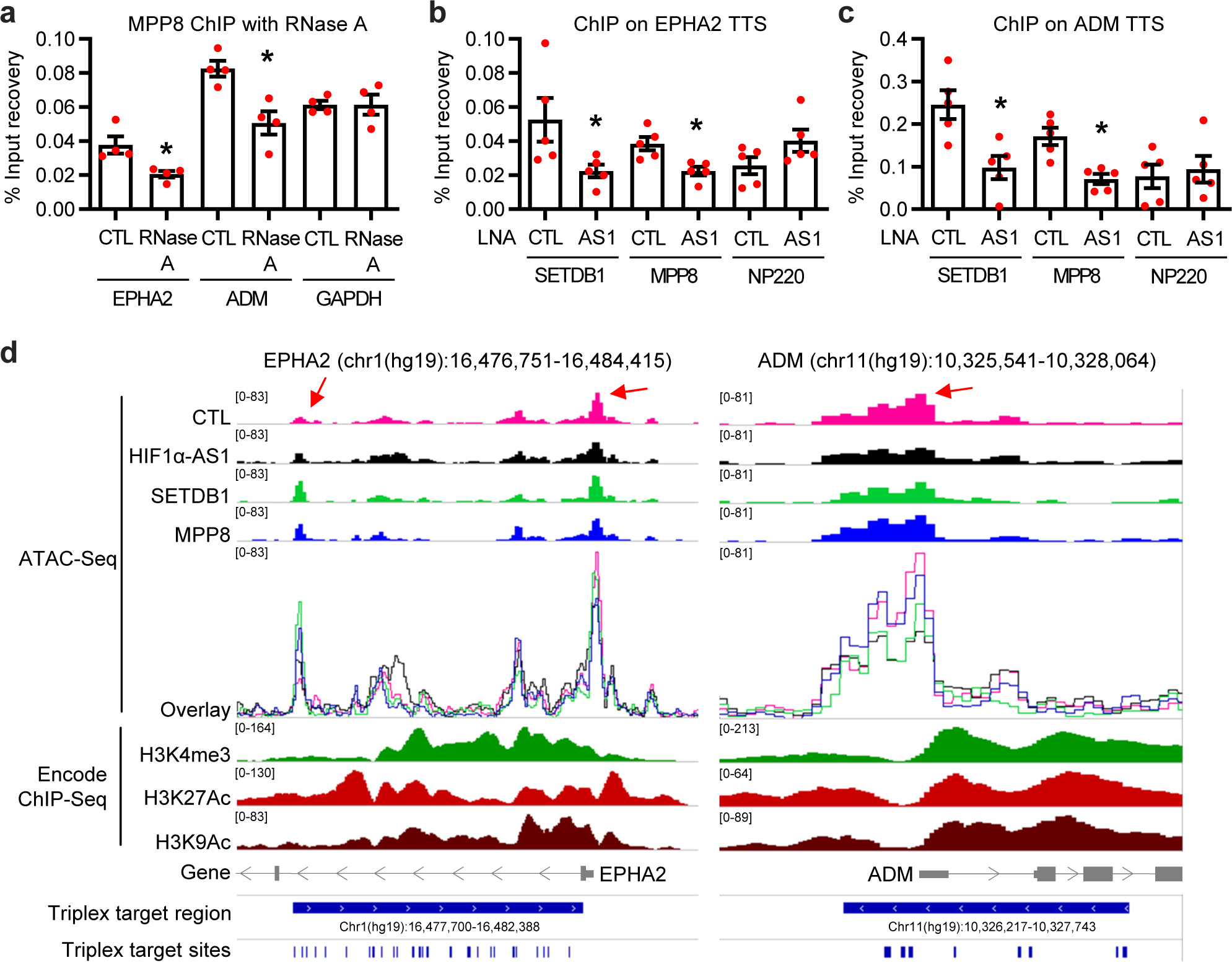
HIF1α-AS1 directs the HUSH complex member MPP8 and SETDB1 to triplex target sites. **a**, Chromatin immunoprecipitation (ChIP) with MPP8 antibodies with or without RNase A treatment and qPCR for the triplex target sites of EPHA2 and ADM. Primers against a promoter sequence of GAPDH served as negative control. n=4, paired t-test. **b-c**, ChIP with antibodies against SETDB1, MPP8 or NP220 in HUVECs treated with (AS1) or without (CTL) LNA GapmeRs against HIF1α-AS1. QPCR was performed for EPHA2 TTS (b) or ADM TTS (c). n=5, paired t-test. **d**, IGV original traces loaded of ATAC- Seq in HUVECs separately and as an overlay after knockdown of HIF1α-AS1 (black), SETDB1 (green), MPP8 (blue) or the negative control (pink). ChIP-Seq data (H3K4me3, H3K27Ac, H3K9Ac) in HUVECs was derived from Encode. Numbers in square brackets indicate data range values. Red arrows indicate altered chromatin accessible regions after knockdown. Error bars are defined as mean +/- SEM. *p<0.05.

## Discussion

The present study combined molecular biology, bioinformatics, physiology and structural analysis to identify and establish the lncRNA HIF1α-AS1 as a triplex-forming lncRNA in human endothelial cells. Through *trans*-acting triplex formation by a specific region within HIF1α-AS1, EPHA2 and ADM DNA target sites are primed for their interaction with the HUSH complex members MPP8 and SETDB1 to mediate gene repression through control of chromatin accessibility. Physiologically, the anti-angiogenic lncRNA HIF1α-AS1 is dysregulated in hypoxia and severe angiogenic and pulmonary diseases like CTEPH, IPAH and GBM. Thus, the present work establishes a putative link of a disease-relevant lncRNA and the HUSH complex by triplex formation resulting in the inhibition of endothelial gene expression.

The interaction of chromatin modifying complexes with lncRNAs suggests that lncRNAs have targeting or scaffolding functions within these complexes with the purpose of modulating chromatin structure and thereby regulating gene expression. Most of these lncRNAs have been identified to interact with complexes such as PRC2, SWI/SNF, E2F1 and p300, e.g. MEG3^4^, FENDRR^12^, MANTIS^34^ and KHPS1^7, 10^. In the present work, we identified other silencing complexes that can be targeted by lncRNAs: We demonstrated that HIF1α-AS1 interacts with proteins of the HUSH complex, which mediates gene silencing. HUSH is also involved in silencing extrachromosomal retroviral DNA^35^. Recently it has been shown that the HUSH complex, particularly MPP8, which is downregulated in many cancer types and whose depletion caused overexpression of long interspersed element-1 (LINE-1s) and Long Terminal Repeats, controls type I Interferon signaling involving a mechanism with dsRNA sensing by MDA5 and RIG-I.^36^ Here we report a direct interaction of the HUSH complex members MPP8 and NP220 with HIF1α-AS1. Moreover, we identified Exon1 of HIF1α-AS1 as being critical for this function. It remains unclear whether the complex exists in its published form in endothelial cells. Our data propose that, in endothelial cells, the HUSH complex interacts with H3K9me3 and DNA and that SETDB1 and MPP8, but not NP220, repress gene expression of HIF1α- AS1-specific target genes.

We propose that HIF1α-AS1 mediates the anti-angiogenic effects through triplex-formation with the receptor tyrosine kinase EPHA2 and the preprohormone ADM genes. EPHA2 is a major regulator of angiogenic processes since EphA2-deficient mice displayed impaired angiogenesis in response to ephrin-A1 stimulation *in vivo*^37^. EphA2-deficient endothelial cells failed to undergo cell migration and vascular assembly in response to ephrin-A1 and only adenovirus-mediated transduction of EPHA2 restored the defect^37^. Additionally, the preprohormone ADM promotes arterio- and angiogenesis^29^. Both genes were upregulated after HIF1α-AS1 knockdown, explaining why HIF1α-AS1 knockdown increased sprouting. However, other HIF1α-AS1 targets are likely to contribute to the phenotype, such as the proangiogenic genes HIF1α^38^, THBS1^39^, EGR1^40^ or NR2F2^41^.

In our unbiased approach, a large number of DNA binding sites were identified for HIF1α-AS1 with triplex domain finder analysis. The large number is not unusual as many of these binding sites overlap and are not identical. Also for other lncRNAs, such as GATA6-AS, FENDRR, HOTAIR and PARTICLE, many DNA binding sites have been predicted within their target genes^9, 14^. EPHA2 and ADM, as well as PLEC, RP11-276H7.2, MIDN and EGR1 contained a large number of DNA binding sites for HIF1α-AS1 and were upregulated after HIF1α-AS1 knockdown. It is therefore tempting to speculate that similar regulatory mechanisms may play a role in the regulation of these genes. For the other target genes, no expression regulation could be found, raising the possibility that DNA binding of HIF1α-AS1 could also have unknown effects such as on splicing or the regulation of binding to promoter elements, histones, transcription factors or 3D chromatin structures.

The evidence for triplex formation by HIF1α-AS1 is based on a number of findings: Firstly, target recognition by HIF1α-AS1 occurs via triplex formation involving GA-rich sequences of the DNA targets and GA-rich sequences within HIF1α-AS1 lncRNA. This has also been observed for other lncRNAs such as HOTAIR^42^ and MEG3^4^, albeit without using RNAs with different TFR lengths, as was the case here for HIF1α-AS1 (27 nt, 46 nt, 131 nt). Secondly, the ^1^H-1D NMR and CD spectroscopy data for HIF1α- AS1 provided similar but more detailed characteristics for triplex formation, compared with other studies^4, 5^. Through the use of heteroduplex samples, measurements at different temperatures, a reduction of equivalents of RNA and triplex analysis with stabilized DNA hairpin sequences, our study allowed an improved and extended analysis of triplex formation. Thirdly, in agreement with previous work^5^, most of the triplex target sites were located in the promoter region or introns of the DNA target genes. Fourthly, the triplex formation of HIF1α-AS1 resulted in gene repression, a finding also observed for other triplex forming RNAs^3^. We could extend this finding by replacing the TFR2 of HIF1α-AS1 with other sequences, which abolished the repressive effects.

HIF1α-AS1 was downregulated in the lungs of patients with specific forms of pulmonary arterial hypertension (PAH). PAH is characterized by several structural changes, remodelling and lesion development in the pulmonary arteries. A study by Masri *et al*. demonstrated the impairment of pulmonary artery endothelial cells from IPAH patients to form tube-like structures^43^. CTEPH, a complex disorder with major vessel remodeling and small vessel arteriopathy, is characterized by medial hypertrophy, microthrombi formation and plexiform lesions^44^. It has been further shown that TGF-ß-induced angiogenesis was increased by circulating CTEPH microparticles co-cultured with pulmonary endothelial cells, indicating a pro-angiogenic feedback of endothelial injury^45^. Since HIF1α-AS1 knockdown led to an increase in sprouting, we assume that the loss of HIF1α-AS1 is a compensatory mechanism, which could be putatively included in the above mentioned pro-angiogenic feedback loop. HIF1α-AS1 was also reduced in endothelial cells isolated from glioblastoma. Typically this pathology represents a highly angiogenic situation with defective endothelium and abnormal morphology^46^. Additionally, HIF1α-AS1 is pro-apoptotic^26^ and so the reduction of HIF1α-AS1 could explain the observed sprouting phenotype by the inhibition of apoptosis. Therefore, it is tempting to speculate that HIF1α-AS1 harbors atheroprotective roles, which could be exploited to alter angiogenesis in patients. Strategies to design such therapeutics require data in other species and in different tissues. HIF1α-AS1 is not endothelial-specific according to CAGE analysis. A comprehensive analysis on HIF1α-AS1 conservation, especially of TFR2, is lacking. Initial attempts with BLAT showed that the first 1000 nt of the pre-processed HIF1α-AS1 including TFR2 were conserved in primates and pigs, but not in rodents (data not shown).

Additionally, the data indicates that triplex formation could have therapeutic potential. The single nucleotide polymorphism (SNP) rs5002 (chr11:10326521 (hg19)) was found within the triplex target site of ADM with phenoscanner, which lists an association with hemoglobin concentration, red blood cell count and hematocrit^47^. Another link between a triplex forming lncRNA and PAH was reported by a massive upregulation of MEG3 in paSMCs from IPAH patients. This prevented hyperproliferation after MEG3 knockdown and a reduced apoptosis phenotype of IPAH-paSMCs involving a mechanism with miR-328-3p and IGF1R^48^. Although triplex formation was not studied, another study provided evidence that a ribonucleotide sequence can be used to form a potential triple helix to inhibit gene expression of the IGF1R gene in rat glioblastoma cells^49^. MEG3 is known to impair cell proliferation and to promote apoptosis in glioma cells^50^. This argues that the binding of a lncRNA to DNA is potentially involved in PAH and GBM.

Taken together, the findings presented here highlight a novel pathway of a scaffolding lncRNA within an epigenetic-silencer complex that has a crucial role in the regulation of endothelial genes.

## Online Methods

### Materials

The following chemicals and concentrations were used for stimulation: Human recombinant VEGF-A 165 (R&D, 293-VE), Recombinant Human FGF-basic (154 a.a.) (bFGF, Peprotech, 100-18B), RNase A (NEB, EN0531) and RNase H (NEB, M0297L). The following antibodies were used: Anti-beta-actin (Sigma-Aldrich, A1978), Anti-H3-pan (Diagenode, C15200011), Anti-dsDNA [35I9 DNA] (Abcam, ab27156), Anti-DNA-RNA Hybrid [S9.6] (Kerafast, ENH001), Anti-EPHA2 (Bethyl, A302-025-M), Anti-GAPDH (Sigma, G8795), Anti-HSC70/HSP70 (Enzo Life Sciences, ADI-SPA-820), Anti-MPP8 (Bethyl, A303-051A-M), Anti-H3K9me3 (Diagenode, SN-146-100), Anti-SETDB1 (Bethyl, A300-121A, for chromatin immunoprecipitation; Santa Cruz Biotechnology, ESET (G-4): sc-271488, for Proximity ligation assay) and Anti-ZNF638/NP220 (Bethyl, A301-548A-M).

### Cell culture

Pooled human umbilical vein endothelial cells (HUVECs) were purchased from Lonza (CC-2519, Lot No. 371074, 369146, 314457, 192485, 186864, 171772, Walkersville, MD, USA). HUVECs were cultured in a humidified atmosphere of 5% CO2 at 37 °C. Fibronectin-coated (356009, Corning Incorporated, USA) dishes were used to culture the cells. Endothelial growth medium (EGM), consisting of endothelial basal medium (EBM) supplemented with human recombinant epidermal growth factor (EGF), EndoCGS-Heparin (PeloBiotech, Germany), 8% fetal calf serum (FCS) (S0113, Biochrom, Germany), penicillin (50 U/mL) and streptomycin (50 µg/mL) (15140-122, Gibco/ Lifetechnologies, USA) was used. For each experiment, at least three different batches of HUVEC from passage 3 were used. In case of hypoxic treatments, cells were incubated in a SciTive Workstation (Baker Ruskinn, Leeds, UK) at 0.1% O2 and 5% CO2 for the times indicated.

### Analyses of Triplex-Seq data to identify candidate lncRNAs

Triplex-Seq data of U2OS and HeLa S3 was used from ^15^, aligned using STAR^51^ and peak-calling performed with MACS2^52^. Peaks were intersected with Ensembl hg38 gene coordinates to produce a list of gene-associated peaks, which was filtered for lncRNAs. The overlap of U2OS and HeLa S3 lncRNAs was filtered for high confidence candidates by applying cut-off filters for fold enrichment (>10) and -log10(P) (>20). Next, the candidates were filtered for the presence of a nuclear value (> 0) in Encode and for the presence of a signal (> 0) in aorta, artery, lymphatic, microvascular, thoracic, umbilical vein and vein in FANTOM5 CAGE data^19–21^. Subsequently, the remaining candidates (RMRP, HIF1α-AS1, RP5-857K21.4, SCARNA2 and SNHG8) were tested for their non-coding probability with the online tools CPAT^53^ and CPC2^54^. Lastly, regions enriched in the Triplex-Seq were manually inspected in the IGV browser to rule out the possibility that the signals belong to overlapping genes.

### Total and nuclear RNA isolation, Reverse transcription and RT-qPCR

Total RNA isolation was performed with the RNA Mini Kit (Bio&Sell). Reverse transcription was performed with SuperScript III Reverse Transcriptase (Thermo Fisher) and oligo(dT)23 together with random hexamer primers (Sigma). CopyDNA amplification was measured with RT-qPCR using ITaq Universal SYBR Green Supermix and ROX as reference dye (Bio-Rad, 1725125) in an AriaMX cycler (Agilent). Relative expression of target genes was normalized to ß-Actin or 18S ribosomal RNA. Expression levels were analyzed by the delta-delta Ct method with the AriaMX qPCR software (Agilent). Oligonucleotides used for amplification are listed in table 1.

For nuclear RNA isolation, cells were resuspended in buffer A1 (10 mM HEPES pH 7.6, 10 mM KCl, 0.1 mM EDTA pH 8.0, 0.1 mM EGTA pH 8.0, 1 mM DTT, 40 µg/mL PMSF) and incubated on ice for 15 min. Nonidet was added to a final concentration of 0.75% and cells were centrifuged (1 min, 4 °C, 16,000 g). The pellet was washed twice in buffer A1, lysed in buffer C1 (20 mM HEPES pH 7.6, 400 mM NaCl, 1 mM EDTA pH 8.0, 1 mM EGTA pH 8.0, 1 mM DTT, 40 µg/mL PMSF) and centrifuged (5 min, 4 °C, 16,000 g). The supernatant was used for RNA isolation with RNA Isolation the RNA Mini Kit (Bio&Sell).

### Knockdown procedures

For small interfering RNA (siRNA) treatments, endothelial cells (80–90% confluent) were transfected with GeneTrans II according to the instructions provided by MoBiTec (Göttingen, Germany). The following siRNAs were used: siEPHA2 (Thermo Fisher Scientific, HSS176396), siSETDB1 (Thermo Fisher Scientific, s19112) and siMPP8 (Thermo Fisher Scientific, HSS123184). As negative control, scrambled Stealth RNAi™ Med GC (Life technologies) was used. All siRNA experiments were performed for 48 h.

For Locked nucleic acid (LNA)-GapmeR (Exiqon) treatment, the transfection was performed with the Lipofectamine RNAiMAX (Invitrogen) transfection reagent according to manufacturer’s protocol. All LNA-GapmeR transfections were performed for 48 h. LNA-GapmeRs were designed with the Exiqon LNA probe designer and contained the following sequences: HIF1α-AS1 (1) 5’-GAAAGAGCAAGGAAC A-3’ and as a negative Control 5’-AACACGTCTATACGC-3’.

### Protein Isolation and Western Analyses

HUVECs were washed in Hanks solution (Applichem) and afterwards lysed with Triton X-100 buffer (20 mM Tris/HCl pH 7.5, 150 mM NaCl, 10 mM NaPPi, 20 mM NaF, 1% Triton, 2 mM Orthovanadat (OV), 10 nM Okadaic Acid, protein-inhibitor mix (PIM), 40 µg/mL Phenylmethylsulfonylfluorid (PMSF)). The cells were centrifuged (10 min, 16,000 g) and protein concentration of the supernatant was determined with the Bradford assay. The cell extract was boiled in Laemmli buffer and equal amounts of protein were separated with SDS-PAGE. The gels were blotted onto a nitrocellulose membrane and blocked in Rotiblock (Carl Roth, Germany). After incubation with the first antibody, infrared-fluorescent-dye-conjugated secondary antibodies (Licor, Bad Homburg, Germany) were used and signals detected with an infrared-based laser scanning detection system (Odyssey Classic, Licor, Bad Homburg, Germany).

### Human Lung samples

The study protocol for tissue donation from human idiopathic pulmonary hypertension patients was approved by the ethics committee (Ethik Kommission am Fachbereich Humanmedizin der Justus Liebig Universität Giessen) of the University Hospital Giessen (Giessen, Germany) in accordance with national law and with Good Clinical Practice/International Conference on Harmonisation guidelines. Written informed consent was obtained from each individual patient or the patient’s next of kin (AZ 31/93, 10/06, 58/15).^55^

Human explanted lung tissues from subjects with IPAH, CTEPH or control donors were obtained during lung transplantation. Samples of donor lung tissue were taken from the lung that was not transplanted. All lungs were reviewed for pathology and the IPAH lungs were classified as grade III or IV.

### PASMC isolation and culture

Pulmonary arterial smooth muscle cells (PASMCs) were handled and treated as described before^56^. Briefly, segments of PASMCs, which were derived from human pulmonary arteries (<2 mm in diameter) of patients with IPAH or from control donors, were cut to expose them to the luminal surface. Gentle scraping with a scalpel blade was used to remove the endothelium. The media was peeled away from the underlying adventitial layer. 1-2 mm^2^ sections of medial explants were cultured in Promocell smooth Muscle Cell Growth Medium 2 (Promocell, Heidelberg, Germany). For each experiment, cells from passage 4-6 were used. A primary culture of human PASMCs was obtained from Lonza (CC-2581, Basel, Switzerland), grown in SmGM-2 Bulletkit medium (Lonza) and cultured in a humidified atmosphere of 5% CO2 at 37 °C. Cells from passages 4-6 were used for experiments. For hypoxia experiments, PASMCs were incubated in hypoxia or normoxia chambers for 24 h in hypoxic medium (basal medium containing 1% FCS for human PASMCs). Hypoxia chambers were equilibrated with a water-saturated gas mixture of 1% O2, 5% CO2, and 94% N2 at 37 °C.

### Brain microvessel isolation from glioblastoma (GBM) patients

Human Brain microvessel (HMBV) isolation from GBM patients was performed exactly as described before.^34^

### CRISPR/dCas9 activation (CRISPRa) and inactivation (CRISPRi)

Guide RNAs (gRNA) were designed with the help of the web-interfaces of CRISPR design (http://crispr.mit.edu/). CRISPR activation (CRISPRa) was performed with a catalytically inactive Cas9 (dCas9), which is fused to the transcription activator VP64 (pHAGE EF1α dCas9-VP64), whereas CRISPRi was performed with a dCas9 fusion to the KRAB repressive domain. Both were used together with a sgRNA(MS2) vector containing the individual guide RNA (gRNA) to induce or repress HIF1α- AS1 gene expression. pHAGE EF1α dCas9-VP64 and pHAGE EF1α dCas9-KRAB were a gift from Rene Maehr and Scot Wolfe (Addgene plasmid # 50918, # 50919)^57^ and sgRNA(MS2) cloning backbone was a gift from Feng Zhang (Addgene plasmid # 61424)^58^. The following oligonucleotides were used for cloning of the guide RNAs into the sgRNA(MS2) vector: For CRISPRa of HIF1α-AS1 5’-CACCGGGGC CGGCCTCGGCGTTAAT-3’ and 5’-AAACATTAACGCCGAGGCCGGCCCC-3’, and for CRISPRi of HIF1α-AS1 5’-CACCGGTCTGGTGAGGATCGCATGA-3’ and 5’-AAACTCATGCGATCCTCACCAGACC-3’. After cloning, plasmids were purified and sequenced. The transfection of the plasmids in HUVEC was performed using the NEON electroporation system (Invitrogen).

### CRISPR-Cas9 genome editing

For genome editing, the ArciTect Cas9-eGFP system was used according to the manufacturer’s conditions (STEMCELL Technologies, Köln, Germany). Briefly, ArciTect™ CRISPR-Cas9 RNP Complex solution was generated with 60 μM gRNA and tracrRNA and 3.6 µg ArciTect™ Cas9-eGFP Nuclease. Afterwards, 20 µM single-strand oligodeoxynucleotide (ssODN) was added to the RNP solution. The following gRNA was used to target TFR2 of HIF1α-AS1: 5‘-ACGTGCTCGTCTGTGTTTAG-3‘. The following ssODNs (Integrated DNA Technologies, Leuven, Belgium) were used to replace TFR2: MEG3, 5‘-GAGGCACAGCTGGGACGGGCTGCGACGCTCACGTGCTCGTCTGTGTTGTAATCGCTCCCTCTCTGCTCTCCGATGGGGGTGCGGCTCAGCCCGAGTCTGGGGACTCTGCGCCTTCTCCGAAGGAA GGCGG-3‘, negative control Luc 5‘-GCTGAGGCACAGCTGGGACGGGCTGCGACGCTCACGTGCTCGTCTGTGTTGTAATTATCACGCTCGTCGTTCGGTATGATGGGGGTGCGGCTCAGCCCGAGTCTGGGGACTCTGCGCCTTCTCCGAAGGAAG-3‘. 400.000 HUVECs were seeded in a 12-well plate and electroporated in E2 buffer with the NEON electroporation system (Invitrogen) (1,400 V, 1x 30 ms pulse). A full medium exchange was done every 24 h and cells were incubated for 72 h.

### HIF1α-AS1 mutants and pCMV6-MPP8-10xHis

To clone pcDNA3.1+HIF1α-AS1, HIF1α-AS1 was amplified with PCR from cDNA (forward primer: 5’- ATATTAGGTACCCGCCGCCGGCGCCCTCCATGGTG-3’, reverse primer: 5’-ACGGGAATTCTAATGGAACATTTCTTCTCCCTAG-3’) and insert and vector (pcDNA3.1+) were digested with Acc65I/EcoRI and ligated. pCMV6-MPP8-MYC-DDK was obtained from Origene (#RC202562L3).

To create pcDNA3.1+HIF1-AS1-Δexon1 (1-116), pcDNA3.1+HIF1-AS1-Δexon2 (117-652), pcDNA3.1+HIF1-AS1-Δexon1 (26-78) and pCMV6-MPP8-10xHIS (replacement of c-terminally MYC-DDK by 10xHIS), site-directed mutagenesis was performed with the Q5 Site-Directed Mutagenesis Kit (NEB) according to the instructions of the manufacturer. Oligonucleotides and annealing temperatures for mutagenesis were calculated with the NEBaseChanger online tool from NEB. The pcDNA3.1+HIF1α-AS1 and pCMV6-MPP8-Myc-DDK plasmids served as templates and were amplified with PCR with the following oligonucleotides to obtain the individual constructs: for pcDNA3.1+HIF1α-AS1-Δexon1 (1-116), 5’-ACTACAGTTCAACTGTCAATTG-3’ and 5’-GGTACCAAGCTTAAGTTTAAAC-3’, for pcDNA3.1+HIF1-AS1-Δexon2 (117-652), 5’- GAATTCTGCAGATATCCAG-3’ and 5’-CTTTCCTTCTCTTCTCCG-3’, for pcDNA3.1+HIF1α-AS1-Δexon1 (26- 78), 5’-AGCGCTGGCTCCCTCCAC-3’ and 5’-TTCACCATGGAGGGCGCC-3’, for pCMV6-MPP8-10xHIS, 5’- CACCATCATCACCACCATCACTAAACGGCCGGCCGCGGTCAT-3’ and 5’- GTGATGGTGAGAGCCTCCACCCCCCTGCAGCTGCACTCTGTATGCACCTATTAGC-3’. The plasmids were verified by sequencing.

To generate purified MPP8-10xHIS protein, pCMV6-MPP8-10xHIS was overexpressed in HEK293 with Lipofectamine 2000 according to the manufacturer’s protocol. Cells were lysed with three cycles snap freezing in nitrogen and 2% triton X-100 with protease inhibitors. Recombinant MPP8-10xHis was purified using HisTrap FF crude columns (Cytiva Europe, Freiburg, Germany, #11000458) with a linear gradient of imidazole (from 20 to 500 mM, Merck, Burlington, United States, #104716) in an Äkta Prime Plus FPLC system (GE Healthcare/Cytiva Europe).

### In vitro transcription and RNA 3’end biotinylation

Prior to *in vitro* transcription, pcDNA3.1+HIF1α-AS1, pcDNA3.1+HIF1α-AS1-Δexon1 (1-116), pcDNA3.1+HIF1α-AS1-Δexon2 (117-652), pcDNA3.1+HIF1α-AS1-Δexon1 (26-78) or control pcDNA3.1+ were linearized with SmaI (Thermo Fisher, FD0663). After precipitation and purification of linearized DNA, DNA was *in vitro* transcribed according to the manufacturers protocol with T7 Phage RNA Polymerase (NEB), and DNA was digested with RQ DNase I (Promega). The remaining RNA was purified with the RNeasy Mini Kit (Qiagen) and used for binding reactions with MPP8-10xHis in RIP experiments. For RNA pulldown experiments, RNA of HIF1α-AS1 or of the control pcDNA3.1+ were further biotinylated at the 3’end with the Pierce RNA 3’end biotinylation kit (Thermo Fisher).

### RNA pulldown assay and mass spectrometry

The RNA pulldown assay was performed similar to^34^. For proper RNA secondary structure formation, 150 ng of 3’end biotinylated HIF1α-AS1 or control RNA was heated for 2 min at 90 °C in RNA folding buffer (10 mM Tris pH 7.0, 0.1 M KCl, 10 mM MgCl2), and then put on RT for 20 min. 1x10^7^ HUVECs were used per sample. Isolation of nuclei was performed with the truCHIP™ Chromatin Shearing Kit (Covaris, USA) according to the manufacturers protocol without shearing the samples. Folded Bait RNA was incubated in nuclear cell extracts for 3 h at 4 °C. After incubation, samples were UV crosslinked. Afterwards, Streptavidin M-270 Dynabeads (80 µL Slurry, Thermo Fisher) were incubated with cell complexes for 2 h at 4 °C. After 4 washing steps with the lysis buffer of the truCHIP chromatin Shearing Kit (Covaris, USA), beads were put into a new Eppendorf tube. For RNA analysis, RNA was extracted with TRIzol (Thermo Fisher). Afterwards, RNA purification was performed with the RNeasy Mini Kit (Qiagen). If indicated, RT-qPCR was performed. For mass spectrometric measurements in order to reduce complexity, samples were eluted stepwise from the beads.

Method description and mass spectrometry proteomics data have been deposited to the ProteomeXchange Consortium (http://proteomecentral.proteomexchange.org) via the PRIDE partner repository^59^ with the dataset identifier PXD023512. Therefore the samples were labelled H1-H5 for HIF1α-AS1 and C1-C5 for the negative control RNA.

### RNA immunoprecipitation

1x10^7^ HUVECs were used per sample. Nuclei isolation was performed with the truCHIP™ Chromatin Shearing Kit (Covaris, USA) according to the manufacturers protocol without shearing the samples. After pre-clearing with 20 µL DiaMag Protein A and Protein G (Diagenode), 10% of the pre-cleared sample served as input and the lysed nuclei were incubated with the indicated antibody or IgG alone for 12 h at 4 °C. The complexes were then incubated with 50 µL DiaMag Protein A and Protein G (Diagenode) beads for 3 h at 4 °C, followed by 4 washing steps in Lysis Buffer from the truCHIP™ Chromatin Shearing Kit (Covaris, USA). In case of RNase treatments, the samples were washed once in TE-buffer and then incubated for 30 min at 37 °C in buffer consisting of 50 mM Tris-HCl pH 7.5-8.0, 150 mM NaCl, 1 mM MgCl2 containing 2 µL RNase H per 100 µL buffer. Afterwards the samples were washed in dilution buffer (20 mmol/L Tris/HCl pH 7.4, 100 mmol/L NaCl, 2 mmol/L EDTA, 0.5% Triton X-100, 1 µL Superase In (per 100 µL) and protease inhibitors). Prior to elution, beads were put into a new Eppendorf tube. RNA was extracted with TRIzol (Thermo Fisher) followed by RNA purification with the RNeasy Mini Kit (Qiagen), reverse transcription and qRT-PCR.

For the *in vitro* RIP assay, the individual RNAs were folded as mentioned above in RNA folding buffer (10 mM Tris pH 7.0, 0.1 M KCl, 10 mM MgCl2), and then put on RT for 20 min. The binding reaction with purified MPP8-10xHIS was performed for 2 h at 4 °C in binding buffer (20 mmol/L Tris/HCl pH8.0, 150 mmol/L KCl, 2 mmol/L EDTA pH 8.0, 5 mmol/L MgCl2, 2 µL/mL Superase In and protease inhibitors). After pre-clearing with 20 µL DiaMag Protein A and Protein G (Diagenode), 5% of the pre-cleared sample served as input. The mixture was incubated with an MPP8 antibody for 3 h at 4 °C.

The complexes were then incubated with 50 µL DiaMag Protein A and Protein G (Diagenode) beads for 1 h at 4 °C, followed by 4 washing steps (5 min, 4 °C, each) in binding buffer. Elution, RNA extraction and RT-qPCR were performed as mentioned above. RT-qPCR was performed with primers targeting the MCS within the in vitro transcribed sequences before (5’-GTGCTGGATATC TGCAGAATTC-3’) and after (5’-GTGCTGGATATCTGCAGAATTC-3’) the HIF1α-AS1 sequences.

### Assay for Transposase Accessibility (ATAC)-Sequencing

ATAC-Seq was performed similar to^34^. 100.000 HUVECs were used for ATAC library preparation using Tn5 Transposase from Nextera DNA Sample Preparation Kit (Illumina). Cell pellets were resuspended in 50 µL PBS and mixed with 25 µL TD-Buffer, 2.5 µL Tn5, 0.5 µL 10% NP-40 and 22 µL H2O. The mixture was incubated at 37 °C for 30 min followed by 30 min at 50 °C together with 500 mM EDTA pH 8.0 for optimal recovery of digested DNA fragments. 100 µL of 50 mM MgCl2 was added for neutralization. The DNA fragments were purified with the MinElute PCR Purification Kit (Qiagen). Amplification of library together with indexing was performed as described elsewhere^60^. Libraries were mixed in equimolar ratios and sequenced on NextSeq500 platform using V2 chemistry and assessed for quality by FastQC. Reaper version 13-100 was employed to trim reads after a quality drop below a mean of Q20 in a window of 5 nt^61^. Only reads above 15 nt were cleared for further analyses. These were mapped versus the hg19 version of the human genome with STAR 2.5.2b using only unique alignments to exclude reads with uncertain arrangement. Reads were further deduplicated using Picard 2.6.0 (Picard: A set of tools (in Java)^62^ for working with next generation sequencing data in the BAM format) to avoid PCR artefacts leading to multiple copies of the same original fragment. The Macs2 peak caller (version 2.1.0)^52^ as employed in punctate mode to accommodate for the range of peak widths typically expected for ATAC-seq. The minimum qvalue was set to -4 and FDR was changed to 0.0001. Peaks overlapping ENCODE blacklisted regions (known misassemblies, satellite repeats) were excluded. Peaks were annotated with the promoter (TSS +/- 5000 nt) of the gene most closely located to the centre of the peak based on reference data from GENCODE v19. To compare peaks in different samples, significant peaks were overlapped and unified to represent identical regions. The counts per unified peak per sample were computed with BigWigAverageOverBed (UCSC Genome Browser Utilities, http://hgdownload.cse.ucsc.edu/downloads.html). Raw counts for unified peaks were submitted to DESeq2 (version 1.14.1) for normalization^63^. Spearman correlations were produced to identify the degree of reproducibility between samples using R. To permit a normalized display of samples in IGV, the raw BAM files were normalized for sequencing depth (number of mapped deduplicated reads per sample) and noise level (number of reads inside peaks versus number of reads not inside peaks). Two factors were computed and applied to the original BAM files using bedtools genomecov resulting in normalized BigWig files.

For samples used after siRNA-mediated silencing of MPP8 and SETDB1 as well as the corresponding LNA GapmeR knockdown of HIF1α-AS1, the improved OMNI-ATAC protocol^64^ was used and samples were sequenced on a Nextseq2000. The resulting data were trimmed and mapped using Bowtie2^65^. Data were further processed using deepTools^66^. For visualization, the Integrative Genomics Viewer^67^ was used.

### Electrophoretic mobility shift assay (EMSA)

RNA transcripts corresponding to HIF1α-AS1 TFR2 region were produced by *in vitro* transcription using the MEGAscript T7 Transcription Kit (Invitrogen) with DNA templates containing the T7 promoter and the sequence to be transcribed. The 131 nt template was produced by PCR using genomic DNA and sequence specific primers, of which the forward one contains the T7 promoter as extention. The DNA templates for the 27 nt and 46 nt transcripts were created by hybridization of single stranded oligos (Sigma) creating a partially (at the T7 promoter sequence) double-stranded molecule.

Triplex target DNA was created by hybridization of equimolar concentrations of short complementary DNA oligos corresponding to the target region in question, whereby only the purine-rich one was ^32^P-γATP-end labelled using T4 PNK enzyme and cleaned with Ethanol precipitation to remove unincorporated hot ATP. This strategy avoids visualization of any RNA:DNA hybrids, that may occur between single stranded molecules. The two oligos were then heated to 70 °C for 10 min after which gradually decreasing the temperature (0.1 °C/sec) to 20 °C, in a buffer containing 10 mM Tris-acetate pH 7.4, 5 mM MgOAc and 50 mM NaCl.

For triplex formation, different amounts of the respective RNA transcripts (50-250 pmol, as indicated) were incubated in a 10 µL reaction with 0.25 pmol of radiolabeled duplex oligos for 1 h at 37 °C in 40 mM Tris-acetate pH 7.4, 30 mM NaCl, 20 mM KCl, 5 mM MgOAc, 10% glycerol and PhosSTOP EASYpack (Roche). For monitoring of triplex formation, the reactions were loaded on a 12% polyacrylamide-bisacrylamide gel containing 40 mM Tris-Ac pH 7.4 and 5 mM MgOAc and run at 120V for 2-3 h at RT. The gels were subsequently dried and exposed a phosphoimager screen overnight, which was then scanned in Fujifilm BAS 1800-II Phosphoimager using the BAS reader 2.2.6 software. Triplex formation was observed as an RNA-dependent shift of the hot duplex oligo as a result of its binding by the RNA and thus slower migration.

Specific sequences for EMSA design and oligonucleotide preparation are shown in tables 2-4.

**Table 3.**
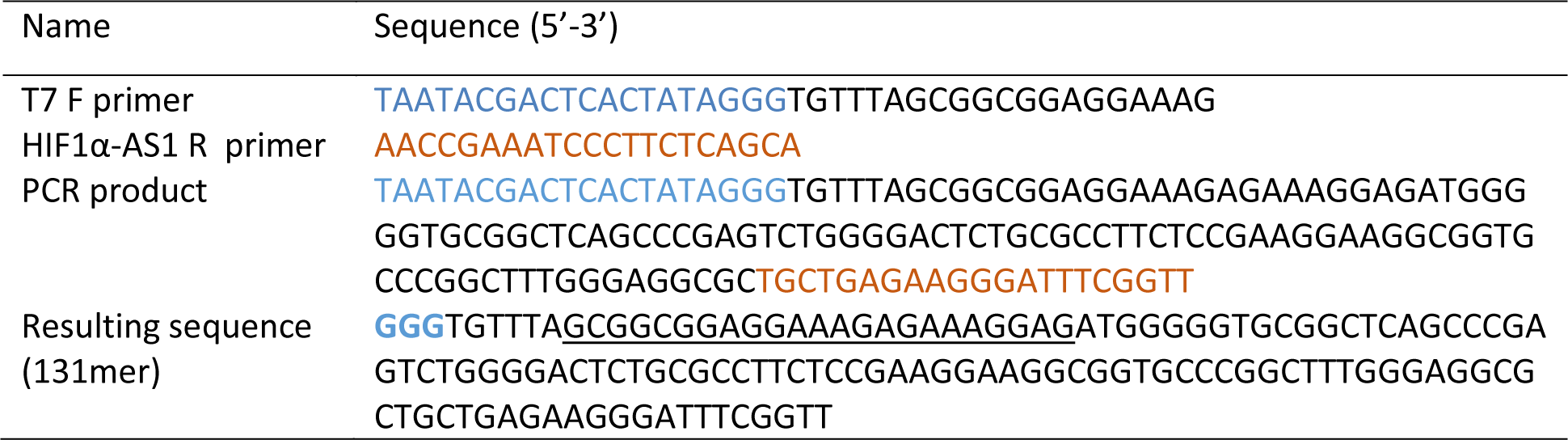
Oligos for generation of the DNA template by PCR for *in vitro* transcription of RNA (131mer).

**Table 4.**
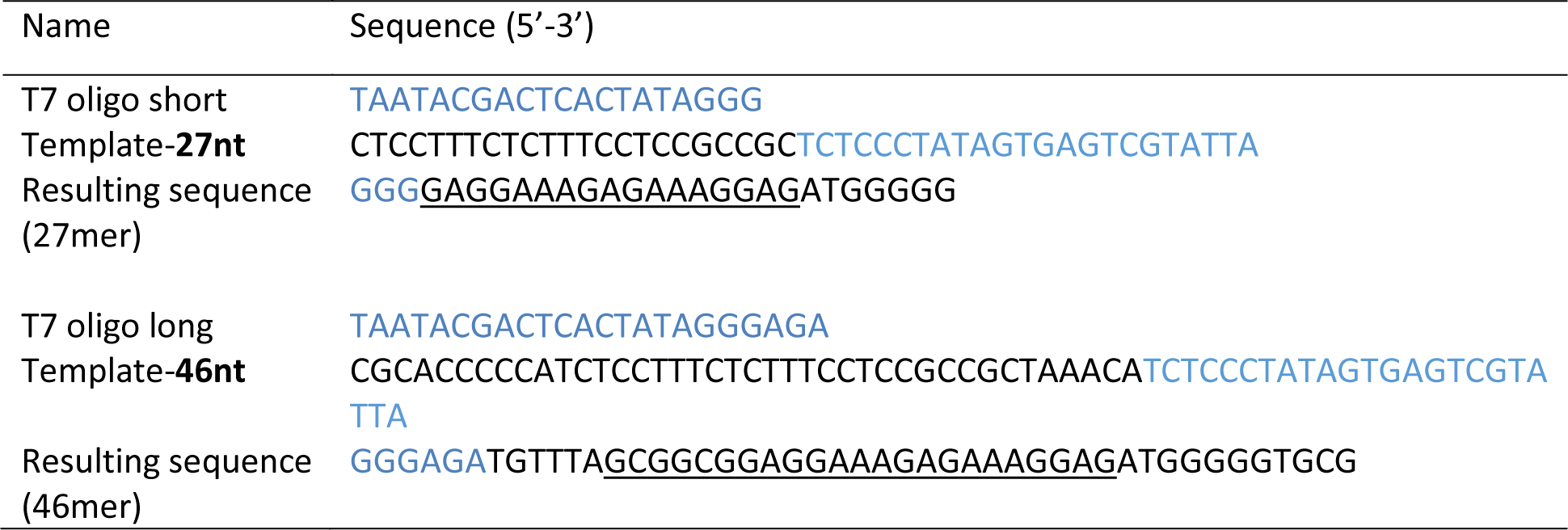
Oligos for generation of partially double stranded DNA template for *in vitro* transcription of RNA (27mer, 46mer).

### RNA and DNA Hybridization

By hybridization of the RNA strand to the DNA duplex or DNA hairpin DNA:DNA:RNA triplexes were formed. First the complementary DNA single strands were incubated at 95 °C for 5 min in hybridization buffer (25 mM HEPES, 50 mM NaCl, 10 mM MgCl2 (pH 7.4)) and afterwards cooled down to RT. Triplex formation was performed by adding RNA to previously hybridized double stranded DNA for 1 h at 60 °C and then cooled down to RT.^13^ For the ^1^H-1D NMR, CD and melting curve experiments, the HIF1α-AS1-TFR2 (TFO2-23) sequence 5‘-GCG GCGGAGGAAAGAGAAAGGAG-3‘ (length 23nt, GC=50.9%) was used in combination with the DNA sequences listed in table 5.

**Table 5.**
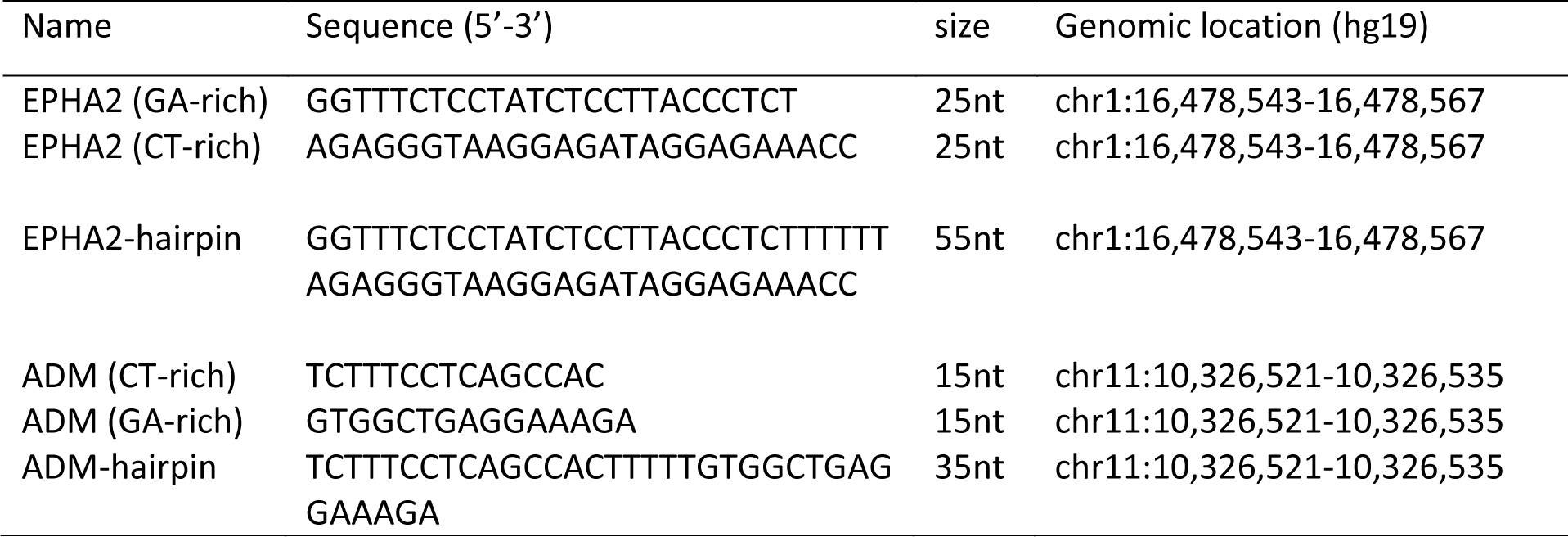
DNA oligos used for ^1^H-1D NMR, CD and melting curve analysis analysis.

### CD spectroscopy and melting curve analysis

Circular dichroism spectra were acquired on a Jasco J-810 spectropolarimeter. The measurements were recorded from 210 to 320 nm at 25 °C using 1 cm path length quartz cuvette. CD spectra were recorded on 8 µM samples of each DNA duplex, DNA:RNA heteroduplex and DNA:DNA:RNA-triplex in 25 mM HEPES, 50 mM NaCl, 10 mM MgCl2 (pH 7.4). Spectra were acquired with 8 scans and the data was smoothed with Savitzky-Golay filters. Observed ellipticities recorded in millidegree (mdeg) were converted to molar ellipticity [θ] = deg x cm^2^ x dmol^-1^. Melting curves were acquired at constant wavelength using a temperature rate of 1 °C/min in a range from 5 °C to 95 °C. All data were evaluated using SigmaPlot 12.5.

### NMR spectroscopy

All NMR samples were prepared in NMR buffer containing 25 mM HEPES-d18, 50 mM NaCl, 10 mM MgCl2 (pH 7.4) with addition of 5 to 10% D2O. All samples were internally referenced with 2,2- dimethyl-2-silapentane-5-sulfonate (DSS). The final NMR sample concentrations ranged between 50 µM to 300 µM. NMR spectra were recorded in a temperature range from 278 K to 308 K on Bruker 600, 800, 900 and 950 MHz spectrometers. 1H NMR spectra were recorded with jump-return-Echo^68^ and gradient-assisted excitation sculpting^69^ for water suppression. NMR data was collected, processed and analyzed using TopSpin 3.6.2 (Bruker).

### Spheroid outgrowth assay

Spheroid outgrowth assays in HUVEC were performed as described in^70^. Stimulation of Spheroids was performed with the indicated amounts of VEGF-A 165 or bFGF for 16 h. Images were generated with an Axiovert135 microscope (Zeiss). Sprout numbers and cumulative sprout lengths were quantified by analysis with the AxioVision software (Zeiss).

### Proximity ligation assay (PLA)

The PLA was performed as described in the manufacturer’s protocol (Duolink II Fluorescence, OLink, Upsalla, Sweden). HUVECs were fixed in phosphate buffered formaldehyde solution (4%), permeabilized with Triton X-100 (0.2%), blocked with serum albumin solution (3%) in phosphate-buffered saline, and incubated overnight with anti-MPP8, anti-dsDNA, anti-SETDB1 or anti-H3K9me3 antibodies. Samples were washed and incubated with the respective PLA-probes for 1 h at 37 °C. After washing, samples were ligated for 30 min (37 °C). After an additional washing step, the amplification with polymerase was performed for 100 min (37 °C). The nuclei were stained using DAPI. Images (with Alexa Fluor, 546 nm) were acquired by confocal microscope (LSM 510, Zeiss).

### Chromatin Immunoprecipitation

Preparation of HUVEC extracts, crosslinking and isolation of nuclei was performed with the truCHIP™ Chromatin Shearing Kit (Covaris, USA) according to the manufacturers protocol. The procedure was similar to ^71^. The lysates were sonified with the Bioruptur Plus (10 cycles, 30 s on, 90 s off, 4 °C; Diagenode, Seraing, Belgium). Cell debris was removed by centrifugation and the lysates were diluted 1:3 in dilution buffer (20 mmol/L Tris/HCl pH 7.4, 100 mmol/L NaCl, 2 mmol/L EDTA, 0.5% Triton X- 100 and protease inhibitors). Pre-clearing was done with DiaMag protein A and protein G coated magnetic beads (Diagenode, Seraing, Belgium) for 1 h at 4 °C. The samples were incubated over night at 4 °C with the antibodies indicated. 5% of the samples served as input. The complexes were collected with 50 µL DiaMag protein A and protein G coated magnetic beads (Diagenode, Seraing, Belgium) for 3 h at 4 °C, washed twice for 5 min with each of the wash buffers 1-3 (Wash Buffer 1: 20 mmol/L Tris/HCl pH 7.4, 150 mmol/L NaCl, 0.1% SDS, 2 mmol/L EDTA, 1% Triton X-100; Wash Buffer 2: 20 mmol/L Tris/HCl pH 7.4, 500 mmol/L NaCl, 2 mmol/L EDTA, 1% Triton X-100; Wash Buffer 3: 10 mmol/L Tris/HCl pH 7.4, 250 mmol/L lithium chloride, 1% Nonidet p-40, 1% sodium deoxycholate, 1 mmol/L EDTA) and finally washed with TE-buffer pH 8.0. In case of RNase treatments, the samples were washed once in TE-buffer and then incubated for 30 min at 37 °C in buffer consisting of 50 mM Tris-HCl pH 7.5-8.0, 150 mM NaCl, 1 mM MgCl2 containing 2 µL RNase H or 2 µL RNase A per 100 µL buffer. Elution of the beads was done with elution buffer (0.1 M NaHCO3, 1% SDS) containing 1x Proteinase K (Diagenode, Seraing, Belgium) and shaking at 600 rpm for 1 h at 55 °C, 1 h at 62 °C and 10 min at 95 °C. After removal of the beads, the eluate was purified with the QiaQuick PCR purification kit (Qiagen, Hilden, Germany) and subjected to qPCR analysis. As a negative control during qPCR, primer for the promoter of GAPDH were used. The primers are listed in table 6.

**Table 6.**
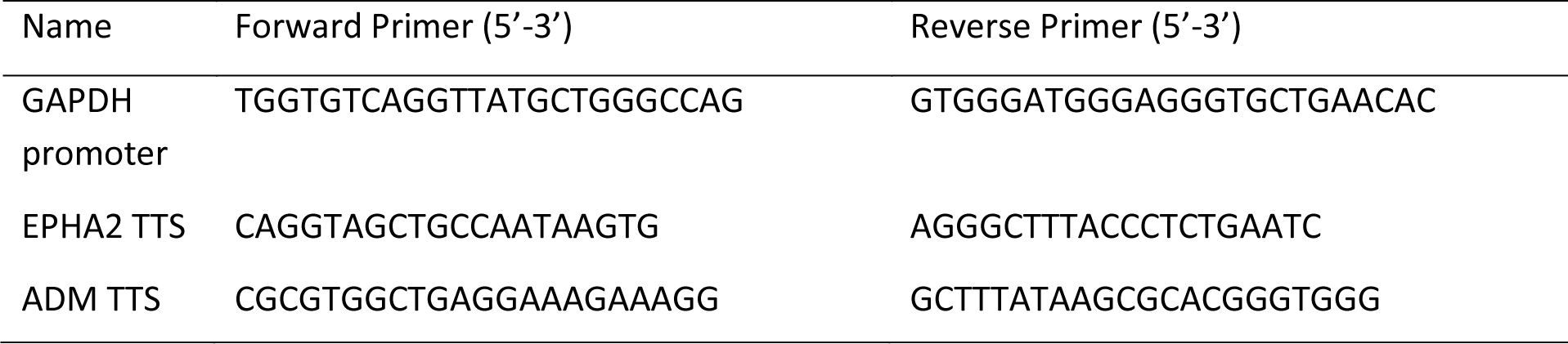
List of primers for ChIP-qPCR.

### Triplex domain finder analysis

Triplex formation of *HIF1α-AS1* was predicted using the Triplex Domain Finder (TDF)^14^ with the human pre-spliced *HIF1α-AS1* sequence (NR_047116.1, gene ID 100750246) to target DNA regions around genes with ATAC-Seq peaks upon HIF1α-AS1 silencing. For annotation of HIF1α-AS1 triplex forming regions across DNA triplex target sites, genome version hg19 was used. Randomization was performed for 200 times. Enrichment was given at a p-value <0.05.

### Data availability

ATAC-Seq data was uploaded to the NCBI SRA database (PRJNA765209, while it remains in private status upon request).

For data about HIF1α-AS1 interaction partners identified with mass spectrometry, the data and methods were uploaded with the dataset identifier PXD023512 to PRIDE (http://www.ebi.ac.uk/pride) and remain in private status upon request.

### Publicly available datasets used

Triplex-Seq data was used from^15^. Fantom5 Encode CAGE expression data was obtained from FANTOM5 website (Gencode v19).^19–21^ ChIP-Seq datasets for HUVEC H3K4me3, H3K27Ac and H3K9Ac were taken from Encode^72^.

### Statistics

Unless otherwise indicated, data are given as means ± standard error of mean (SEM). Calculations were performed with Prism 8.0 or BiAS.10.12. The latter was also used to test for normal distribution and similarity of variance. In case of multiple testing, Bonferroni correction was applied. For multiple group comparisons ANOVA followed by post hoc testing was performed. Individual statistics of dependent samples were performed by paired t-test, of unpaired samples by unpaired t-test and if not normally distributed by Mann-Whitney test. P values of <0.05 was considered as significant. Unless otherwise indicated, n indicates the number of individual experiments.

## Acknowledgments

We thank Cindy F. Höper for excellent technical assistance and Jana Meisterknecht for help with mass spectrometry. We are grateful to Katalin Pálfi and Tanja Lüneburg for the help with cell culture.

This work was supported by the Goethe University Frankfurt am Main, the German Centre for Cardiovascular Research (DZHK, Funding Postdoc start up, Förderkennzeichen: 81X3200107), the DFG excellence cluster Cardiopulmonary Institute (CPI) EXS2026 and the DFG Transregio TRR267 (TP A04, TP A06 and TP Z02). Work at BMRZ is supported by the state of Hesse.

## Author information

These first authors contributed equally: Matthias S. Leisegang, Jasleen Kaur Bains These corresponding authors contributed equally: Harald Schwalbe, Ralf P. Brandes *Author contributions* MSL, JKB, SS, SSP, FR, RG, IW, TR, IGC, HS and RPB designed the experiments. MSL, JKB, SS, JAO, NMK, CCK, SG, CC, FB, JIP, RB, CV, DF, BPM, IW, TR performed the experiments. MSL, JKB, SS, JAO, NMK, NSC, IG, BPM, IW, TR, HS and RPB analyzed the data. SS, JAO, CCK, SG, TW, JP, ML, MHS, RG, IGC performed bioinformatics. NSC, MHS, RG and IG helped with research design and advice. MSL, JKB, HS and RPB wrote the manuscript. All authors interpreted the data and approved the manuscript.

## Supplementary information

**Extended data figure 1:**
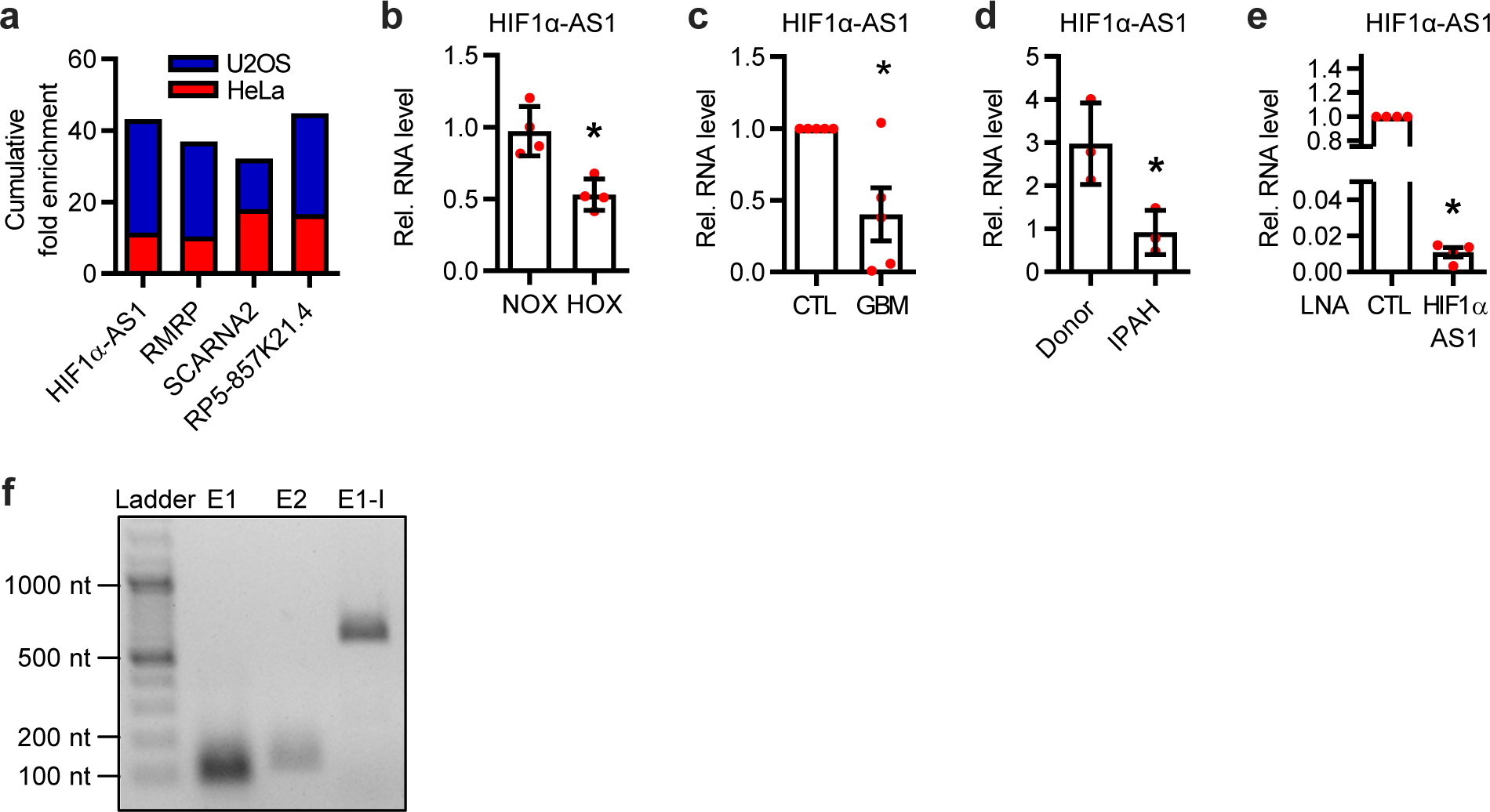
**a**, Cumulative fold enrichment of the four remaining candidates in the U2OS and HeLa S3 Triplex-Seq. **b**, RT-qPCR of HIF1α-AS1 in paSMCs treated under hypoxic conditions (HOX, 1% O2) for 24 h. Cells treated under normoxia (NOX) served as basal control. n=4, Unpaired t-test. **c**, RT-qPCR of HIF1α-AS1 from endothelial cells isolated from glioblastoma (GBM) or adjacent healthy control (CTL) tissue. n=5. Paired t-test. **d**, RT-qPCR of HIF1α-AS1 in paSMCs from control donors (Donor) or patients with idiopathic pulmonary arterial hypertension (IPAH). n=3, Unpaired t-test. **e**, RT-qPCR of HIF1α-AS1 after knockdown with LNA-GapmeRs against HIF1α-AS1 or an LNA negative control (CTL). n=4, Paired t-test. **f**, Agarose gel after RT-PCR of Exon1 (E1), Exon2 (E2) or the first 714nt of the pre-processed HIF1α-AS1 (E1-I). Error bars are defined as mean +/- SEM. *p<0.05.

**Extended data figure 2:**
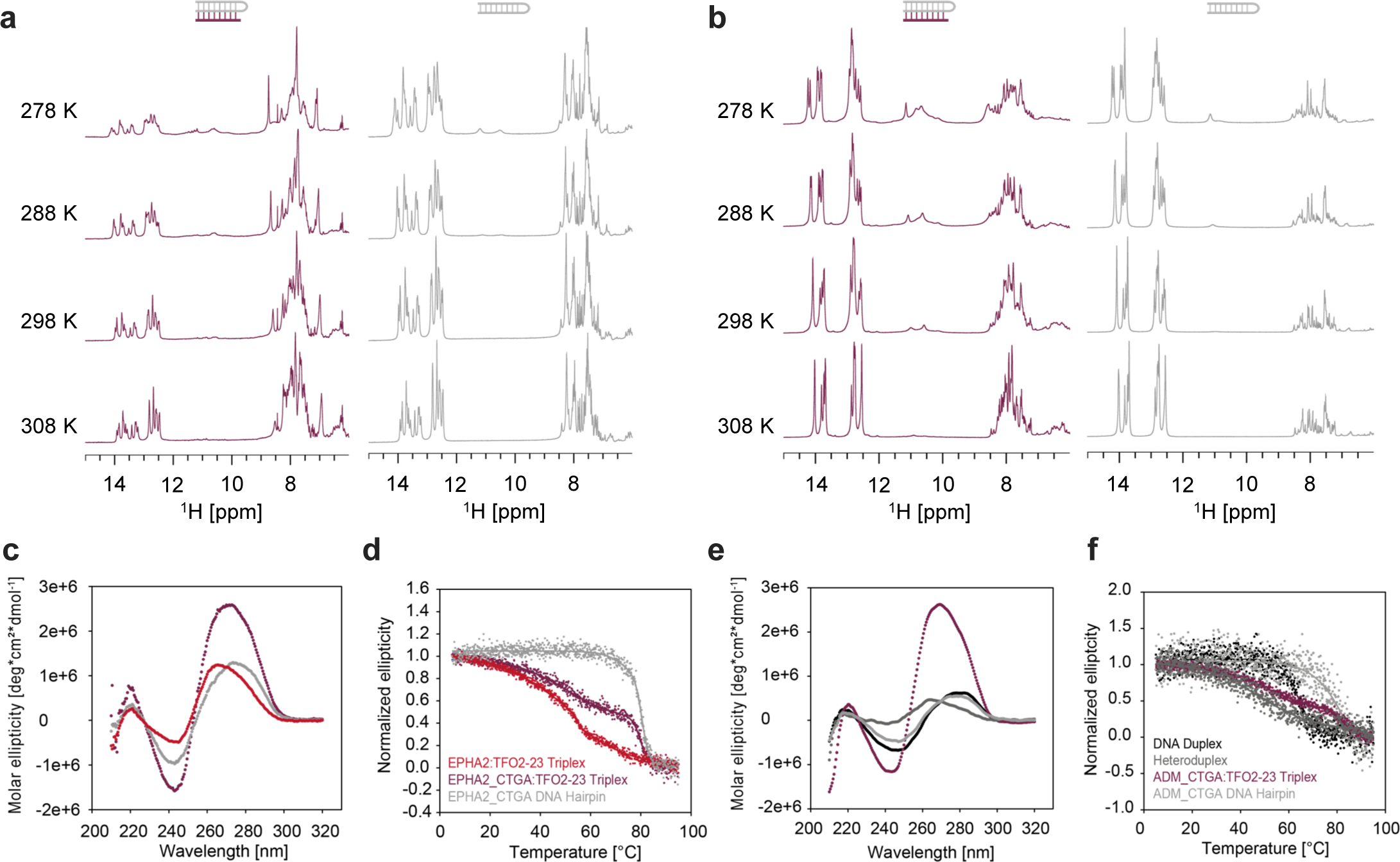
**a**, H-1D NMR spectra of the EPHA2_CTGA hairpin (grey) and the EPHA2_CTGA:HIF1α-AS1-TFR2 triplex (dark red) in a temperature range between 278-308 K. **b**, H-1D NMR spectra of the ADM_CTGA hairpin (grey) and the ADM_CTGA:HIF1α-AS1-TFR2 triplex (dark red) in a temperature range between 278-308 K. **c**, Circular dichroism spectra of the EPHA2:HIF1α-AS1- TFR2 (TFO2-23) triplex (red), the EPHA2_CTGA hairpin alone (light grey) and the EPHA2_CTGA:HIF1α- AS1-TFR2 (TFO2-23) triplex (dark red) measured at 298 K. **d**, UV melting of the EPHA2:HIF1α-AS1- TFR2 (TFO2-23) triplex (red), the EPHA2_CTGA hairpin (light grey) and EPHA2_CTGA:HIF1α-AS1-TFR2 (TFO2-23) (dark red). **e**, Circular dichroism spectra of the the ADM duplex (black), the heteroduplex (dark grey), the ADM_CTGA hairpin alone (light grey) and the ADM_CTGA:HIF1α-AS1-TFR2 (TFO2-23) triplex (dark red) measured at 298 K. **f**, UV melting of the ADM duplex (black), the heteroduplex (dark grey), the ADM_CTGA hairpin (light grey) and ADM_CTGA:HIF1α-AS1-TFR2 (TFO2-23) triplex (dark red).

**Extended data figure 3:**
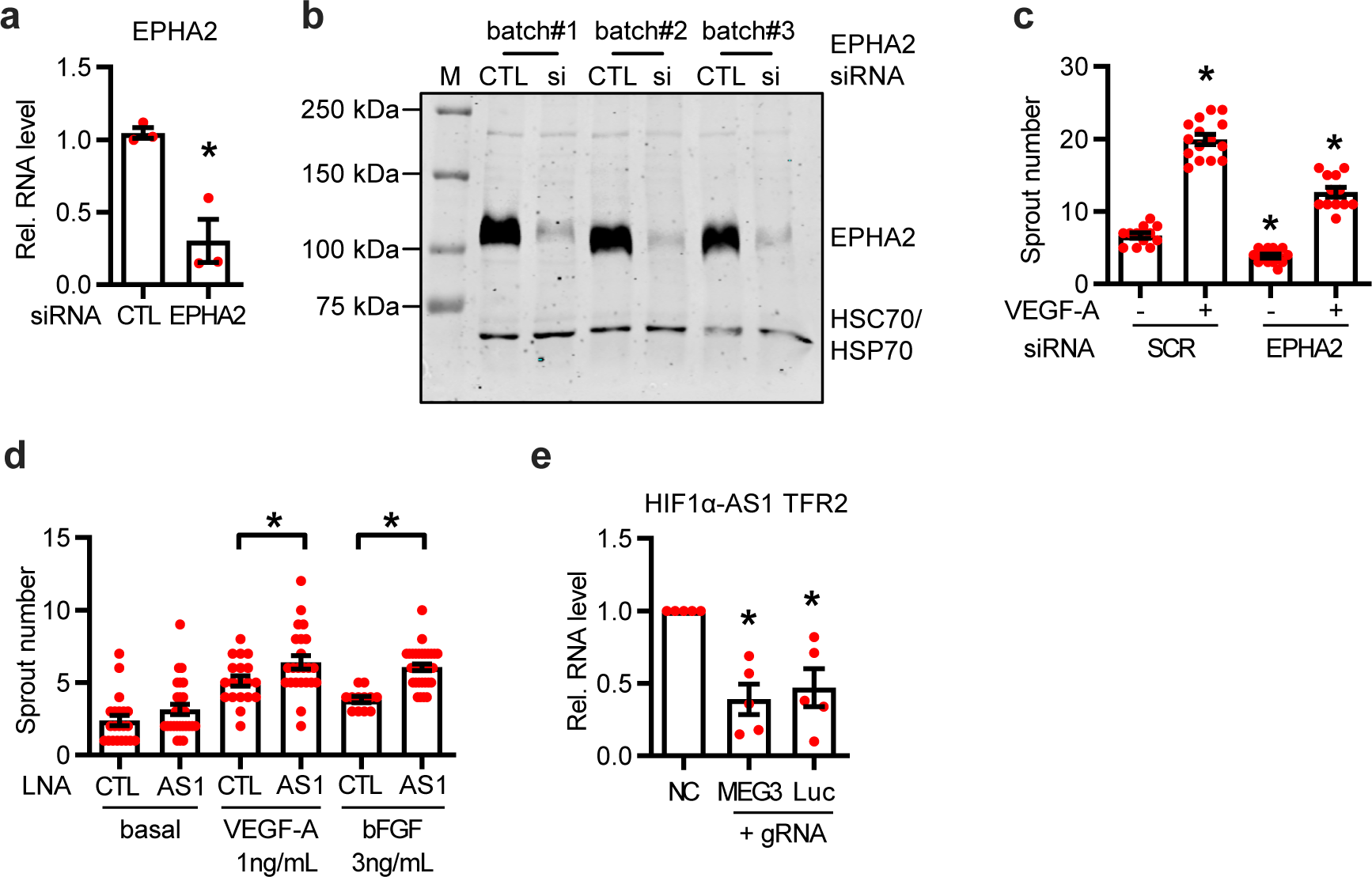
**a**, RT-qPCR after siRNA-mediated knockdown of EPHA2. Expression levels of EPHA2 are shown. Scrambled siRNA (CTL) served as negative control. n=3, Unpaired t-test. **b**, Western blot with (si) or without (CTL) siRNA-mediated knockdown of EPHA2 in three different batches of HUVEC. EPHA2 and HSC70/HSP70 antibodies were used. M, marker. **c**, Quantification of the sprout numbers from the spheroid assay seen in Fig. 4d. One-Way ANOVA with Bonferroni post hoc test. n=12-15. **d**, Quantification of the sprout numbers from the spheroid assay seen in Fig. 4f. One-Way ANOVA with Bonferroni post hoc test. n=12-32. **e**, Relative RNA level of HIF1α-AS1 TFR2 after a ssODN-mediated replacement of the TFR2 within HIF1α-AS1 with the TFR of MEG3 or a DNA fragment of a luciferase negative control. NC, nontemplate control. n=5, Paired t-test. Error bars are defined as mean +/- SEM. *p<0.05.

**Extended data figure 4:**
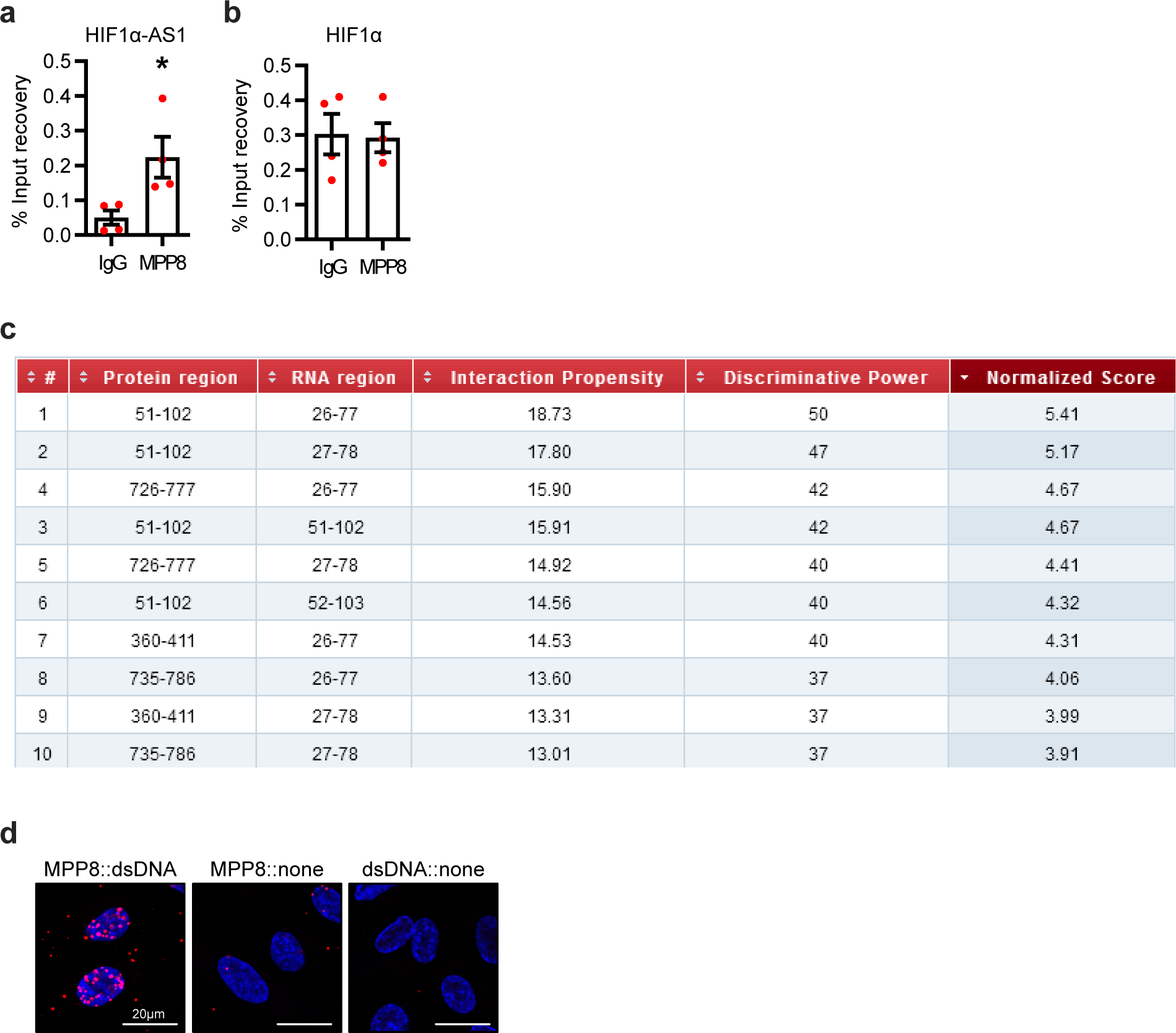
**a&b**, RIP with MPP8 antibodies and qPCR for HIF1α-AS1 (a) or HIF1α (b). IgG served as negative control. n=4, Mann Whitney t-test. **c**, Binding propensity of MPP8 and HIF1α-AS1 calculated with *cat*RAPID. **d**, Proximity ligation assay of HUVECs with antibodies against MPP8 and dsDNA. The individual antibody alone served as negative control. Red dots indicate polymerase amplified interaction signals. Scale bar indicates 20 µm. Error bars are defined as mean +/- SEM. *p<0.05.

Sup. Table 1: Triplex-Seq HeLa S3 lncRNA regions

Sup. Table 2: Triplex-Seq U2OS lncRNA regions

Sup. Table 3: List of TTS of TFR1

Sup. Table 4: List of TTS of TFR2

Sup. Table 5: List of TTS of TFR3

Sup. Table 6: Interaction partners of HIF1α-AS1

